# The Relative Binding Position of Nck and Grb2 Adaptors Dramatically Impacts Actin-Based Motility of Vaccinia Virus

**DOI:** 10.1101/2021.10.07.463509

**Authors:** Angika Basant, Michael Way

## Abstract

Phosphotyrosine (pTyr) motifs in unstructured polypeptides orchestrate important cellular processes by engaging SH2-containing adaptors to nucleate complex signalling networks. The concept of phase separation has recently changed our appreciation of such multivalent networks, however, the role of pTyr motif positioning in their function remains to be explored. We have now explored this parameter in the assembly and operation of the signalling cascade driving actin-based motility and spread of Vaccinia virus. This network involves two pTyr motifs in the viral protein A36 that recruit the adaptors Nck and Grb2 upstream of N-WASP and Arp2/3-mediated actin polymerization. We generated synthetic networks on Vaccinia by manipulating pTyr motifs in A36 and the unrelated p14 from Orthoreovirus. In contrast to predictions, we find that only specific spatial arrangements of Grb2 and Nck binding sites result in robust N-WASP recruitment, Arp2/3 driven actin polymerization and viral spread. Our results suggest that the relative position of pTyr adaptor binding sites is optimised for signal output. This finding may explain why the relative positions of pTyr motifs are usually conserved in proteins from widely different species. It also has important implications for regulation of physiological networks, including those that undergo phase transitions.

## INTRODUCTION

Multicellular animals extensively use phosphotyrosine (pTyr) signals for growth, communication, movement and differentiation (Jin and Pawson, 2012; Lim and Pawson, 2010). Pathways including EGF and insulin receptor signalling, as well as T cell activation rely on pTyr motifs interacting with SH2 domains (Blumenthal and Burkhardt, 2020; Lemmon and Schlessinger, 2010). SH2 domains are present in kinases, phosphatases as well as adaptor proteins that lack enzymatic activity but couple upstream signalling events to downstream function (Liu and Nash, 2012). Examples of such adaptors include Shc1, Crk, Nck and Grb2 that also contain other interaction modules including SH3 domains that bind polyproline motifs (Bywaters and Rivera, 2021; Mayer, 2015). pTyr signalling is often dysregulated in cancers and other diseases. For example, EGF receptors can be mutationally activated or present in high copy numbers vis-à-vis their cognate adaptors, and oncogenic mutations frequently map to SH2 domains (Li et al., 2012a; Li et al., 2021; Shi et al., 2016; Sigismund et al., 2017). It therefore remains important to understand the molecular principles of how pTyr signalling networks function.

pTyr motifs commonly occur in poorly ordered, unstructured regions of proteins (Stavropoulos et al., 2012; Van Roey et al., 2013). Moreover, a single polypeptide may be phosphorylated more than once to generate multiple SH2 domain binding sites creating a large network of interactions with many signalling modules. Indeed, like many receptor tyrosine kinases, the C terminus of EGFR is disordered (Figure 1 – supplement 1A) (Keppel et al., 2017; Pinet et al., 2021). This region contains several pTyr motifs including those that bind Grb2 and Shc1 (Batzer et al., 2006; Lin et al., 2019; Mandiyan et al., 1996; Smith et al., 2006; Ward et al., 1996). The relative positions of these motifs are conserved across vertebrates (Figure 1 – supplement 1B). Though they are short and space-efficient, linear motifs bearing such pTyr must interact with effectors comprising globular domains of varying sizes. Additionally, disordered sequences can become structured when bound to their respective domains (Davey, 2019; Nioche et al., 2002; van der Lee et al., 2013). Given that such factors are likely to impose constraints or “polarity” on the architecture of signalling networks, can the positioning of motifs have an impact on function? If yes, this would suggest that pTyr motifs cannot be repositioned as they are optimised to achieve the desired signalling output.

**Figure 1:**
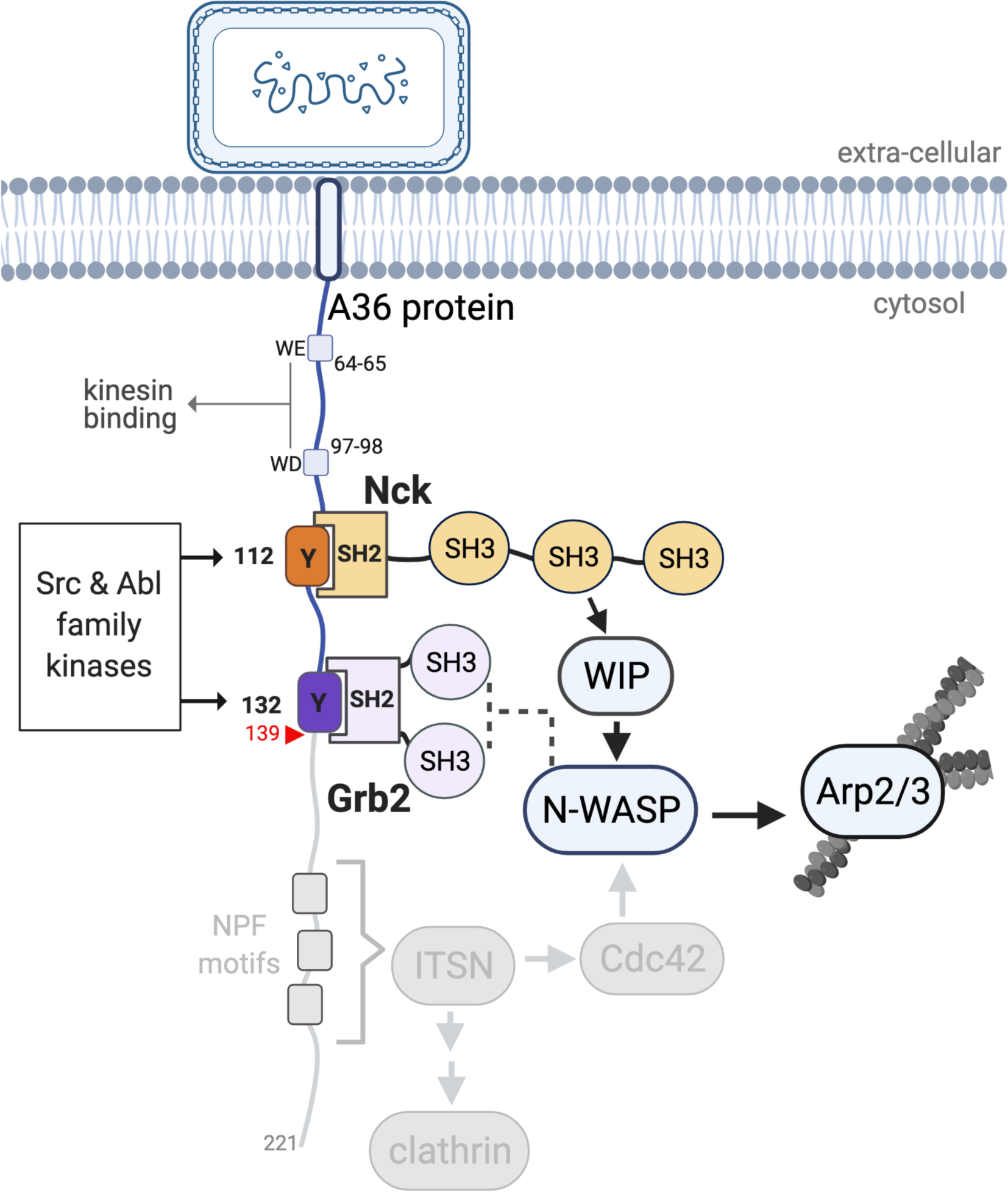
A36 interactions and the Vaccinia signalling network. A schematic showing the Vaccinia virus protein A36 and its known interactors. Kinesin-1 that drives microtubule-based transport of virions to the plasma membrane binds the WD/WE motifs. Nck and Grb2 bind Y112 and Y132 respectively when they are phosphorylated by Src and Abl family kinases. Nck and Grb2 interact with WIP and N-WASP via their SH3 domains, which results in the activation of the Arp2/3 complex and stimulation of actin polymerization. The region deleted in A36 after residue 139 (red triangle) to abolish the involvement of the RhoGEF intersectin and its binding partners clathrin and Cdc42 is shown in grey.

In recent years, the framework of phase separation or biomolecular condensates has shed new light on signalling network organization (Banani et al., 2017; Huang et al., 2019; Li et al., 2012b; Zhao and Zhang, 2020). Integral membrane proteins with disordered cytoplasmic regions and involved in multivalent pTyr-SH2 interactions such as Linker for Activation of T cells (LAT) and the kidney podocyte regulator nephrin form phase separated condensates at critical concentrations (Case et al., 2019; Ditlev et al., 2019; Kim et al., 2019; Pak et al., 2016; Su et al., 2016). This striking property is thought to contribute to their signalling function. As these proteins have been largely investigated by overexpressing components in cells or by reconstitution *in vitro*, we are yet to fully understand how the underlying principles of condensate organisation regulate physiological signalling (Alberti et al., 2019; Mayer and Yu, 2018; McSwiggen et al., 2019). Furthermore, the relationship between pTyr motifs in these systems has only been explored via disruption of the sites by a mutational approach (Huang et al., 2017) and the role of their positioning has never been investigated. Do membrane signalling proteins make stochastic connections with their downstream components or do underlying wiring principles exist? We chose to address the importance of pTyr motif positioning in a physiologically relevant model, that of Vaccinia virus egress from its infected host cell (Leite and Way, 2015).

Following replication and assembly, Vaccinia virus recruits kinesin-1 via WD/WE motifs in the cytoplasmic tail of A36 an integral viral membrane protein to transport virions to the cell periphery on microtubules (Dodding et al., 2011) (Figure 1). After viral fusion with the plasma membrane, extracellular virions that remain attached to the cell locally activate Src and Abl family kinases (Frischknecht et al., 1999; Newsome et al., 2004; Newsome et al., 2006; Reeves et al., 2005). This results in phosphorylation of tyrosine 112 and 132 in a disordered region of A36 once it incorporates into the plasma membrane when the virus fuses at the cell edge (Frischknecht et al., 1999; Newsome et al., 2004; Newsome et al., 2006; Reeves et al., 2005; Ward and Moss, 2004) (Figure 1 & Figure 1 – supplement 1C). pTyr 112 and 132 motifs bind the SH2 domains of adaptors Nck and Grb2 respectively (Scaplehorn et al., 2002). Nck recruits N-WASP via WIP to activate the Arp2/3 complex at the virus (Donnelly et al., 2013). The resulting actin polymerisation drives virus motility and enhances its cell-to-cell spread (Ward and Moss, 2004). Nck is essential for actin tail formation, while Grb2 recruitment helps stabilise the signalling complex (Frischknecht et al., 1999; Scaplehorn et al., 2002; Weisswange et al., 2009). Additionally, NPF motifs in A36 interact with the RhoGEF intersectin to recruit Cdc42 and clathrin to the virus, further enhancing actin polymerisation (Humphries et al., 2012; Humphries et al., 2014; Snetkov et al., 2016) (Figure 1). The turnover rates of Nck, Grb2 and N-WASP beneath extracellular virions are highly reproducible with little variability, suggesting the signalling network has a defined organisation (Weisswange et al., 2009). This underlying signalling network organization may arise from the fact that both WIP and N-WASP only have two Nck-binding sites, each with distinct preferences for the three adaptor SH3 domains (Donnelly et al., 2013). This Nck-dependent signalling network is not unique to Vaccinia and is also used to polymerise actin by nephrin in kidney podocytes and the pathogenic *E. coli* EPEC protein Tir (Hayward et al., 2006; Jones et al., 2006; Welch and Way, 2013).

Cellular pTyr networks are typically transient and can be difficult to detect, making them challenging to manipulate and study quantitatively. In contrast, the Vaccinia virus signalling network and actin tail formation are robust and sustained. Vaccinia therefore provides an *in vivo* genetically tractable system where outputs of actin polymerisation, virus speed and spread can be quantitatively measured. Using recombinant Vaccinia viruses, including ones expressing Orthoreovirus protein p14 (Figure 1 – supplement 1C), we uncovered a striking impairment in actin-based motility and spread of the virus when the relative positions of Nck and Grb2 pTyr binding motifs were exchanged and/or manipulated. Collectively our results indicate that the relative positioning of pTyr motifs and downstream adaptor binding is an important factor in the output of signalling networks that has thus far been overlooked.

## RESULTS

### A minimal Vaccinia system to investigate how pTyr signalling regulates actin polymerization

The recruitment of Nck by A36 at pTyr112 is necessary and sufficient to drive actin-based motility of Vaccinia virus whereas Grb2 recruitment at pY132 improves actin tail formation (Frischknecht et al., 1999; Scaplehorn et al., 2002; Weisswange et al., 2009). In addition to these two adaptors, the recruitment of intersectin, Cdc42 and clathrin by full-length A36 also indirectly regulates the extent of virus-driven actin polymerisation (Humphries et al., 2012; Humphries et al., 2014; Snetkov et al., 2016)2014; Snetkov et al., 2016). To reduce the complexity arising from these interactions and other unknown A36 binding partners, we generated a recombinant virus that recruits a minimal signalling network to activate Arp2/3-dependent actin polymerisation (Figure 1). To achieve this goal, the WR–ΔA36R virus lacking the A36 gene was rescued with an A36 variant that terminates after the Grb2 binding site at residue 139 (referred to as the A36 N-G virus hereafter). This shorter A36 variant retains the WD/WE kinesin-1 binding motifs that are required for microtubule-based transport of the virus to the plasma membrane. As in the full-length protein, actin tail formation induced by this shorter A36 variant is dependent on the Nck-binding site (Figure 2 – supplement 1A, A36 N-G vs X-G). Similar to the full-length protein (Scaplehorn et al., 2002), truncated A36 induces shorter actin tails in the absence of Grb2 recruitment (Figure 2 – supplement 1A, A36 N-G vs N-X). Having confirmed the role of Nck and Grb2 is the same as in the wild-type virus, we used the A36 N-G virus to explore the role of pTyr motif positioning in signalling output.

**Figure 2:**
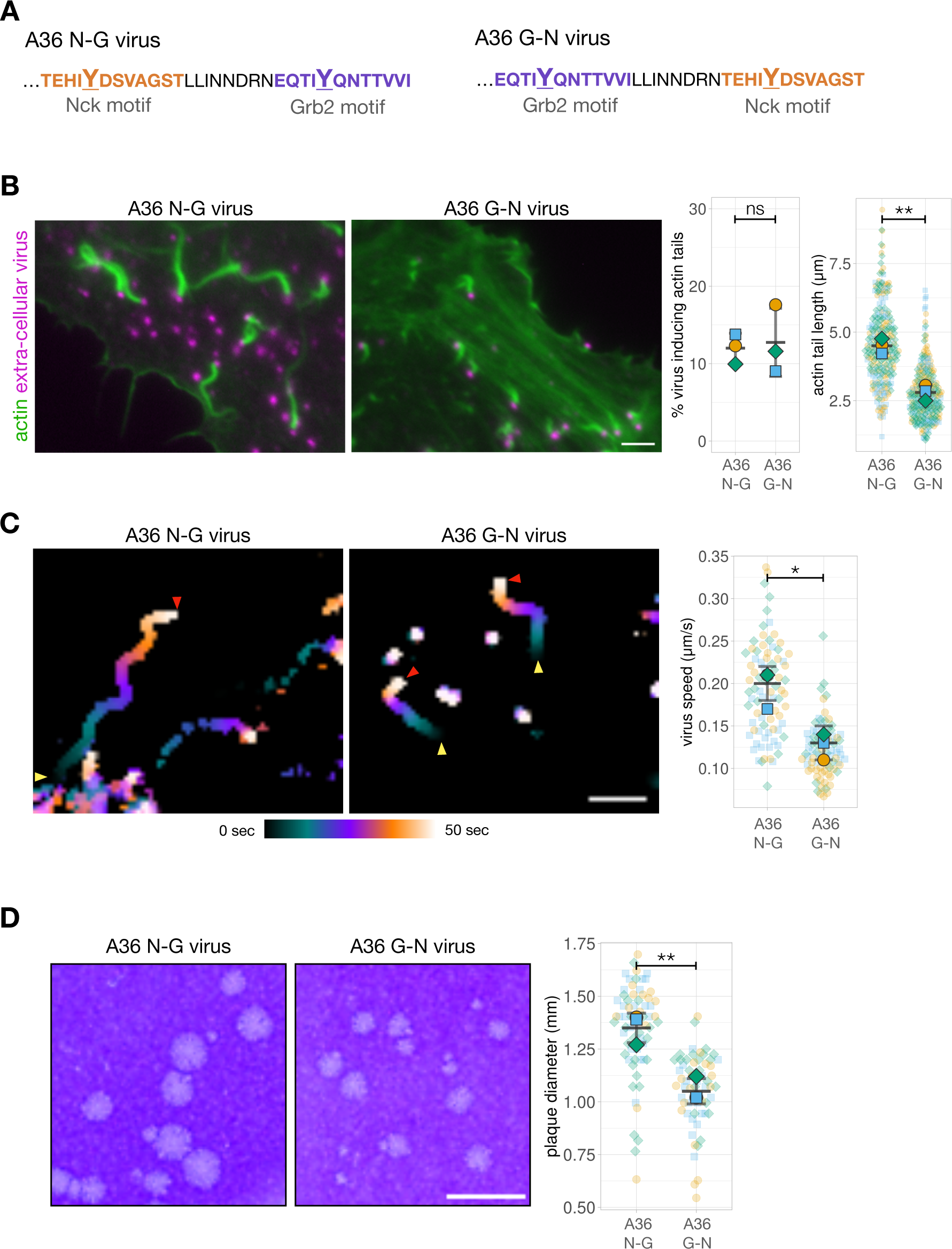
Phosphotyrosine motif position impacts actin-based motility and viral spread. **A.** C-terminal amino acid sequence of A36 in recombinant viruses showing the position of phosphotyrosine motifs in their wild type (A36 N-G) and swapped (A36 G-N) configurations. **B.** Representative immunofluorescence images of actin tails in HeLa cells infected with the indicated virus at 8 hours post-infection. Actin is stained with phalloidin, and extra-cellular virus particles attached to plasma membrane are labelled using an antibody against the viral protein B5. Scale bar = 3 μm. The graphs show quantification of number of extracellular virus particles inducing actin tails and their length. A total of 270 actin tails were measured in three independent experiments. **C.** Temporal colour-coded representation of time-lapse movies tracking the motility of the indicated RFP-A3-labelled virus over 50 seconds at 8 hours post infection (Supplemental Movie 1). Images were recorded every second and the position of virus particles at frame 1 (yellow triangles) and frame 50 (red triangles) are indicated. Scale bar = 3 μm. The graph shows quantification of virus speed over 50 seconds. A total of 82 virus particles were tracked in three independent experiments. **D.** Representative images and quantification of plaque diameter produced by the indicated virus in confluent BS-C-1 cells 72 hours post-infection. 64 plaques were measured in three independent experiments. Scale bar = 3mm. All error bars represent S.D and the distribution of the data from each experiment is shown using a “SuperPlot”. Welch’s t test was used to determine statistical significance; ns, p >0.05; * p ≤0.05; ** p ≤ 0.01.

### The relative positioning of Nck and Grb2-binding has a dramatic impact on actin polymerization

Residues N-terminal to the Tyr (positions -4 to -1) are important for tyrosine kinase site recognition while SH2 domain binding specificity is typically based on +1 to +6 residues C-terminal to the pTyr (Blasutig et al., 2008; Frese et al., 2006; Kefalas et al., 2018; Songyang and Cantley, 1995; Wagner et al., 2013). Given these previous observations, we compared actin polymerisation, virus motility and spread of the A36 N-G virus with a recombinant virus where 12 residues surrounding the pTyr112 Nck-binding site were exchanged with the pTyr132 Grb2 interaction motif (Figure 2A, A36 G-N virus hereafter). Our modified virus thus maintains requirements for Src and Abl mediated phosphorylation as well as Nck and Grb2 SH2 binding. Strikingly, exchanging the positions of these two pTyr motifs has a dramatic impact on actin tail length though the number of extracellular virus particles inducing actin polymerization remains unaltered (Figure 2B). It is possible that this effect is a consequence of the relative positioning of the two adaptor binding sites in A36. Alternatively, it may be that the Nck-binding site is sub-optimal when it is repositioned into the Grb2-binding locus. To determine which is true, we assessed the impact of changing the position of the Nck-binding site in the absence of Grb2 recruitment. We found that viruses A36 N-X and X-N that lack a Grb2 binding site (N-X: Nck binds in native position and X-N: Nck binds in Grb2 position), did not differ in their extent or ability to induce actin tails (Figure 2 – supplement 1B). This indicates that relative positioning of the Nck and Grb2 phosphotyrosine motifs in A36 is important for optimal signalling output (efficient actin polymerization). Actin polymerisation at the virus drives both the motility and spread of virions. To test whether motif positioning had an impact on the former, we imaged cells infected with the A36 N-G or A36 G-N viruses expressing A3, a viral core protein tagged with RFP. In the swapped configuration, corresponding to reduced actin polymerisation, virus motility is slower (Figure 2C, Supplemental Movie 1). In addition, the A36 G-N virus has reduced cell-to-cell spread as it forms smaller plaques on confluent cell monolayers compared to A36 N-G (Figure 2D). Our observations with Vaccinia clearly demonstrate that the output of a signalling network is strongly influenced by the relative positioning of Nck and Grb2 binding sites in the membrane protein responsible for initiating the signalling cascade.

### Inducing actin polymerization using a synthetic signalling network

We were curious whether our observations were unique to Vaccinia A36 and/or if the positioning of adaptor binding sites would also be important in a different context. We took advantage of the Vaccinia signalling platform to generate a synthetic pTyr network that replaces A36 with a different protein capable of activating actin polymerization via the same adaptors. Recent observations demonstrate that p14, an integral membrane protein from Orthoreovirus activates N-WASP via Grb2 to regulate cell fusion (Chan et al., 2020). Grb2 is recruited by a pTyr116 motif located in the short cytoplasmic tail of p14 that is also predicted to be disordered (Figure 1 – supplement 1C). Interestingly, examination of the p14 sequence reveals there are two tyrosine residues located 16 and 20 amino acids upstream of the pTyr116 that are predicted to interact with Nck (Eukaryotic Linear Motif (ELM) and Scansite 4.0) (Kumar et al., 2019; Obenauer et al., 2003) (Figure 3A). To examine whether these sites participate in actin polymerization we generated a hybrid construct comprising the first 105 amino acids of A36 including the kinesin-1-binding motifs required to traffic virions to the plasma membrane and residues 79-125 of p14 (Figure 3A). Transient expression of p14 N-G in cells infected with Vaccinia virus lacking A36 results in extracellular virions inducing robust actin tails that were dependent on the presence of N-WASP (Figure 3B and C). Given this, we mutated Tyr96, 100 and 116 of p14 in turn and examined the impact on actin tail formation. This analysis reveals there is no role for Tyr100 as its mutation to phenylalanine had no impact on actin tail formation (Figure 3D). In contrast, Tyr96 is essential for the virus induced actin polymerization. As with Vaccinia A36, the ability of transiently expressed p14 N-G to promote actin polymerization primarily depends on the predicted Nck-binding pTyr96 and is enhanced by pTyr116, which is a bona fide Grb2 binding site (Chan et al., 2020) (Figure 3D and E). To confirm whether pTyr96 interacts with Nck, we incubated HeLa cell lysates with beads conjugated with peptides containing the prospective binding motif. The pTyr96 peptide retains Nck from the cell lysate but not the unphosphorylated control (Figure 3F). Our observations clearly demonstrate that as with Vaccinia A36, p14 of Orthoreovirus induces actin polymerization using a Nck and Grb2 signalling network. These results also establish that the Vaccinia platform can be used to analyze synthetic signalling networks and test predictions concerning their operation.

**Figure 3:**
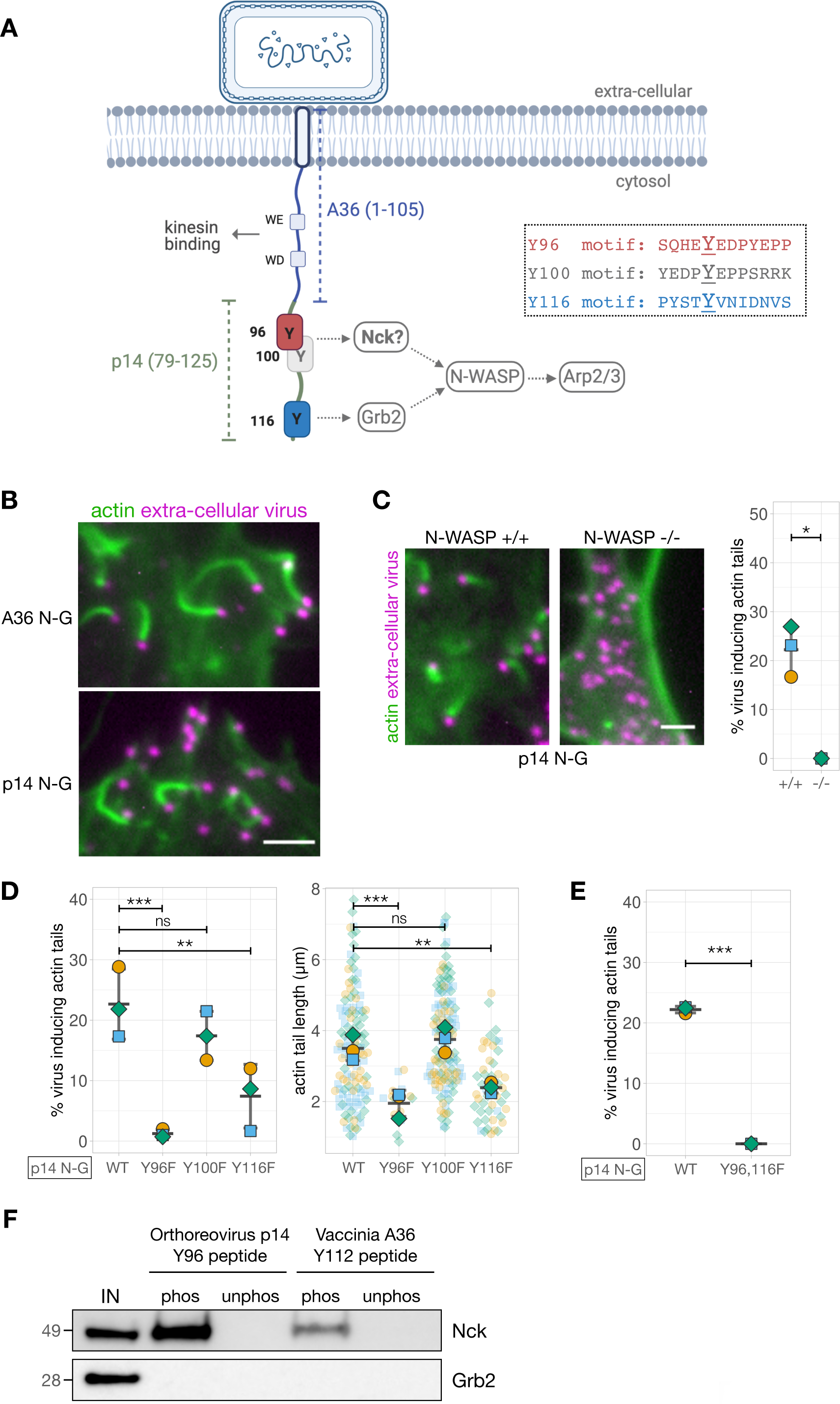
Generating and validating an A36-p14 hybrid that can polymerise actin. **A.** Schematic showing a hybrid construct (referred to as p14 N-G) comprising the first 105 residues of A36 and the C-terminal residues 79-125 of Orthoreovirus p14 protein. Positions of the predicted Nck-binding sites at Tyr96 and Tyr100, and the previously established Grb2- binding site at Tyr116 are shown together with their respective sequences. **B.** Representative immunofluorescence images of actin tails in HeLa cells infected with Vaccinia virus lacking the A36 gene and transiently expressing the indicated constructs under the A36 promoter at 8 hours post-infection. Actin is stained with phalloidin, and extra-cellular virus particles attached to plasma membrane are labelled using an anti-B5 antibody. Scale bar = 3 μm. **C.** Representative immunofluorescence images of N-WASP null or parental mouse embryonic fibroblast cells infected with Vaccinia virus lacking the A36 gene and transiently expressing the p14 N-G construct under the A36 promoter at 16 hours post-infection. Actin is stained with phalloidin, and extra-cellular virus particles are labelled using an anti-B5 antibody. Scale bar = 3 μm. The graph shows quantification of actin tail number per extracellular virus particle. Error bars represent S.D. from three independent experiments. Welch’s t test was used to determine statistical significance; * p ≤0.05. **D** and **E.** Quantification of the number of extracellular virus inducing actin tails together with their length in HeLa cells infected with Vaccinia virus lacking the A36 gene and transiently expressing p14 N-G constructs under the A36 promoter with indicated Tyr to Phe mutations, at 8 hours post-infection. 125 actin tails were measured in three independent experiments, except in mutants Y96F and Y116F where fewer actin tails were made. All error bars represent S.D and the distribution of the data from each experiment is shown using a “SuperPlot”. Dunnett’s multiple comparison’s test (for panel D) and Welch’s t test (for panel E) were used to determine statistical significance; ns, p >0.05; ** p ≤ 0.01; *** p ≤0.001. **F.** Immunoblot analysis of peptide pulldowns showing that endogenous Nck from HeLa cell lysates binds to phosphopeptides corresponding to Tyr96 from the Orthoreovirus p14 and Tyr112 of the Vaccinia A36 but not to their unphosphorylated counterparts.

Given this, we generated a recombinant virus where endogenous A36 was replaced with the hybrid A36-p14 protein (henceforth referred to as the p14 N-G virus). As we have observed with Vaccinia A36, actin tails generated by the p14 N-G virus recruit endogenous Nck, WIP and N-WASP to the virion (Figure 4A). In the absence of an antibody that can detect Grb2, we confirmed the p14 N-G virus also recruits Grb2 by infecting HeLa cells stably expressing GFP-Grb2 (Figure 4A) (Figure 4 – supplement 1). We next sought to determine whether the relative positioning of the Nck and Grb2 binding sites was important for the signalling output of the p14. We therefore generated a recombinant virus (p14 G-N virus) where the Nck- and Grb2-binding motifs of p14 are in a swapped orientation (Figure 4B). Remarkably, as we observed for A36, swapping these motifs did not impact on the number of viruses inducing actin polymerization but resulted in shorter actin tails and a slower virus speed (Figure 4C and D, Supplemental Movie 2). Plaque assays on confluent cell monolayers reveals that the p14 G-N virus is also attenuated in its spread (Figure 4E). Once again, the signalling output of the system clearly depends on the relative positioning of Nck and Grb2 pTyr binding sites.

**Figure 4:**
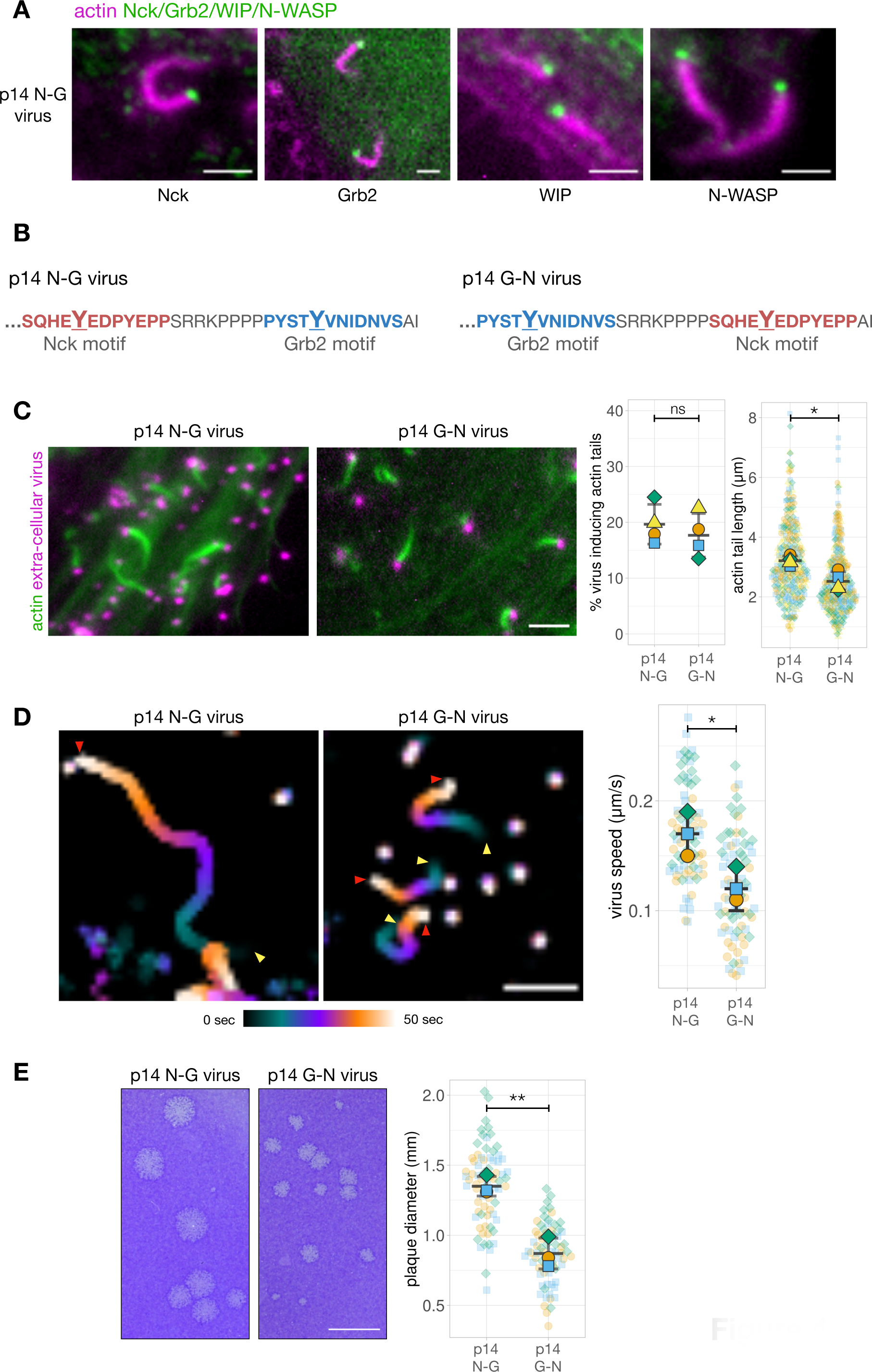
Phosphotyrosine motif position in a A36-p14 hybrid protein impacts actin polymerisation. **A.** Representative images showing the recruitment of Nck, WIP, Grb2 and N-WASP to actin tails in HeLa cells infected with a recombinant virus expressing the p14 N-G construct at the A36 locus at 8 hours post-infection. Endogenous Nck, N-WASP and WIP were detected with antibodies and actin is labelled with phalloidin. To ascertain Grb2 localisation, infected HeLa cells stably expressing GFP-Grb2 and transiently expressing LifeAct iRFP were imaged live. Scale bar = 2 μm. **B.** C-terminal amino acid sequence of the A36 -p14 hybrid in recombinant viruses showing the positions of phosphotyrosine motifs in their wild-type (p14 N-G) and swapped (p14 G-N) configurations. **C.** Representative immunofluorescence images of actin tails in HeLa cells infected with the indicated virus at 8 hours post-infection. Actin is stained with phalloidin, and extra-cellular virus particles are using an anti-B5 antibody. Scale bar = 3 μm. The graphs show quantification of number of extracellular virus particles inducing actin tails and their length. 336 actin tails were measured in four independent experiments. **D.** Temporal colour-coded representation of time-lapse movies tracking the motility of the indicated RFP-A3-labelled virus over 50 seconds at 8 hours post infection (Supplemental Movie 2). Images were recorded every second and the position of virus particles at frame 1 (yellow triangles) and frame 50 (red triangles) are indicated. Scale bar = 3 μm. The graph shows quantification of virus speed over 50 seconds. 75 virus particles were tracked in three independent experiments. **E.** Representative images and quantification of plaque diameter produced by the indicated virus in confluent BS-C-1 cells 72 hours post-infection. 72 plaques were measured in three independent experiments. Scale bar = 3mm. All error bars represent S.D and the distribution of the data from each experiment is shown using a “SuperPlot”. Welch’s t test was used to determine statistical significance; ns, p >0.05; * p ≤0.05; ** p ≤ 0.01.

### Why does the relative position of Nck- and Grb2-binding sites impact actin polymerization?

To further probe into how pTyr motif positioning impacts signalling output, we focused on the A36 signalling network given we have a deeper understanding of how it functions. The simplest explanation for the difference between the signalling output of the N-G and G-N configurations is that the levels of A36 are different between the two viruses. To examine if this is the case, we generated recombinant viruses where the A36 N-G and G-N variants were tagged at their C-terminus with tgGFP2. We measured the fluorescent intensity ratio of tgGFP2 to the viral core protein A3 fused to RFP. RFP-A3 provides a reliable internal reference marker as its fluorescent intensity is consistent across different viruses and cell lines expressing GFP-tagged components of the Vaccinia signalling network (Figure 5 – supplement 1A). We found no significant difference between the levels of A36 N-G and G- N relative to RFP-A3 (Figure 5 – supplement 1B). This suggests that underlying cause for the difference in signalling output resides in the network itself. We therefore determined the level of recruitment of the components involved in activating the Arp2/3 complex on the virus. We infected HeLa cells stably expressing GFP-tagged Nck, Grb2, WIP or N-WASP (Figure 4 – supplement 1) and measured their respective GFP intensities on the virus relative to RFP-A3 as an internal fluorescent standard. We found that the levels of Nck are comparable between the A36 N-G and G-N viruses (Figure 5A). In contrast, Grb2 and N-WASP were significantly reduced on the G-N virus (Figure 5A). The level of WIP, which is also recruited to actin tails is also lower in the G-N configuration (Figure 5 – supplement 1C). To confirm these results, we also took advantage of MEF cell lines stably expressing GFP-tagged Nck- or N-WASP but lacking their respective endogenous proteins (Figure 5 – supplement 2). We found that as seen in HeLa cells, the levels of GFP-Nck were similar for both viruses but the A36 G-N virus recruited two-fold less GFP-N-WASP than the N-G variant (Figure 5B). It is possible that the reduction in N-WASP is a consequence of lower Grb2 levels as this adaptor helps stabilize N-WASP on the virus (Weisswange et al., 2009). To examine this possibility, we measured N-WASP levels on a virus where the Grb2-binding site is abolished by a Tyr132 to Phe mutation (A36 N-X virus). Loss of Grb2 binding had no appreciable impact on the levels of Nck on the A36 N-X virus as compared to those on the A36 N-G and G-N viruses (Figure 5C). The levels of N-WASP on the A36 N-X virus were reduced (46.4%) but not to the same extent as the A36 G-N virus (75%). Repositioning the Nck and Grb2 sites clearly leads to a more severe impairment of the ability of the signalling network to activate Arp2/3 driven actin polymerization than loss of the Grb2 site. Moreover, this suggests that Grb2 has a dominant negative effect when it is mis-positioned relative to Nck.

**Figure 5:**
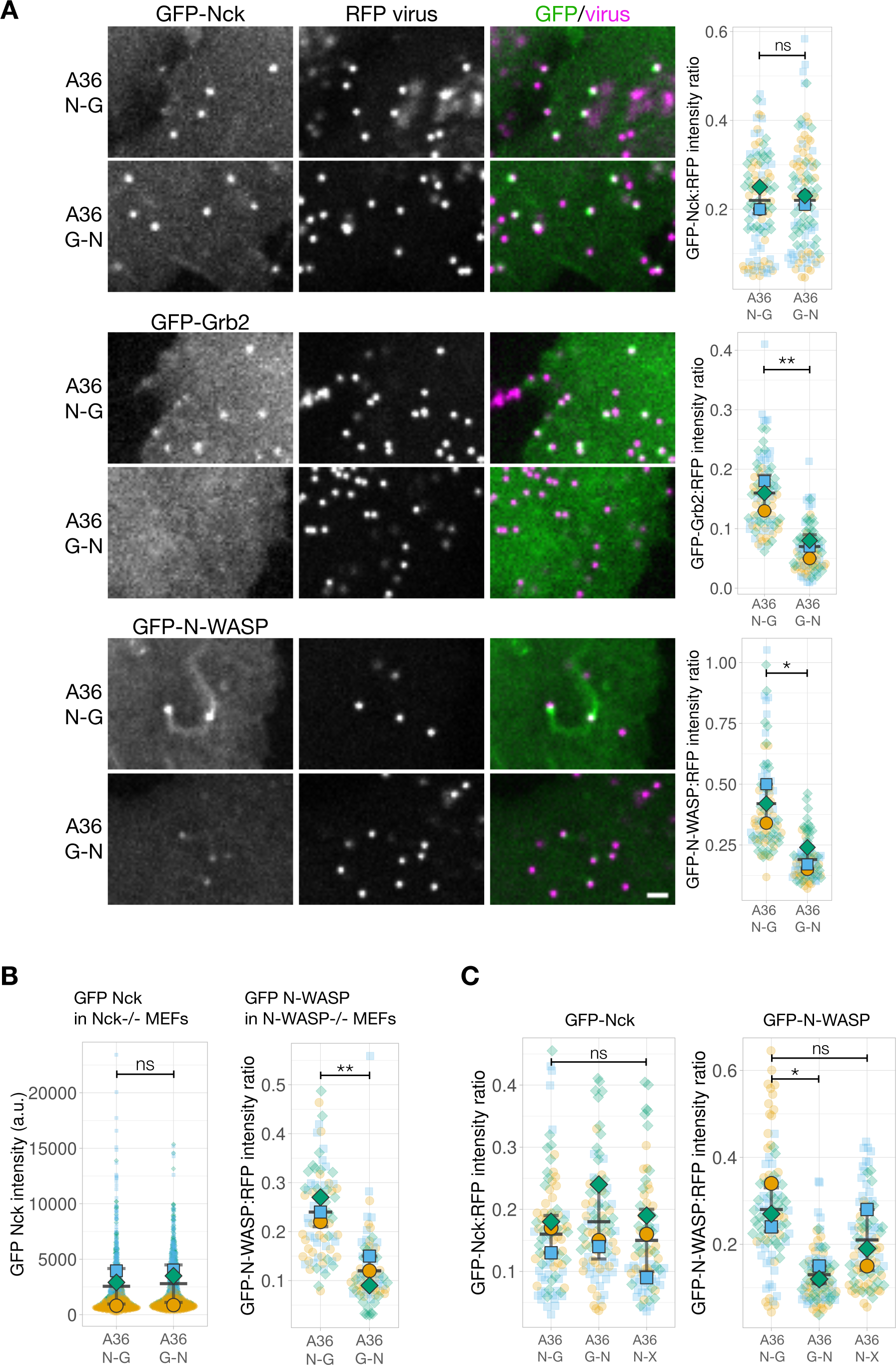
N-WASP recruitment is impaired when Grb2 binding is repositioned in A36. **A.** Representative images showing the indicated GFP-tagged protein recruitment to RFP-A3 labelled virus particles in live HeLa cells infected with the indicated viruses at 8 hours post-infection. Scale bar = 2 μm. The graphs show quantification of GFP:RFP-A3 fluorescence intensity ratio. Intensity of 90 virus particles was measured in three independent experiments. **B.** Left - The graph shows quantification of GFP-Nck intensity at extracellular virus particles in mouse embryonic fibroblasts (MEFs) lacking both Nck1 and Nck2 and stably expressing GFP-Nck1 at 16 hours post-infection. Intensity of 1200 virus particles was measured in three independent experiments. Right - The graph shows quantification of GFP-N-WASP:RFP-A3 fluorescence intensity ratio on virus particles in live N-WASP-/- MEFs stably expressing GFP-N-WASP infected with the indicated viruses at 16 hours post-infection. Intensity of 75 virus particles was measured in three independent experiments. **C.** The graphs show quantification of GFP:RFP-A3 fluorescence intensity ratio on virus particles in live HeLa cells stably expressing the indicated GFP-tagged protein infected with the A36 N-G, G-N or N-X viruses at 8 hours post-infection. Intensity of 90 virus particles was measured in three independent experiments. All error bars represent S.D and the distribution of the data from each experiment is shown using a “SuperPlot”. Dunnett’s multiple comparison’s test (for panel C) or Welch’s t test (remaining panels) were used to determine statistical significance; ns, p >0.05; * p ≤ 0.05; ** p ≤ 0.01.

### Reduced Grb2 recruitment is not due to diminished phosphorylation of Tyrosine 132

The comparable levels of Nck recruitment to the A36 N-G and G-N viruses suggests that phosphorylation of Tyr112 (the Nck-binding site) is not impacted by changing its position within A36. However, as Grb2 recruitment is impaired we sought to determine whether its binding site (Tyr132) is less phosphorylated in the G-N virus. During Vaccinia infection, only a small fraction of the total cellular A36 is phosphorylated by Src and Abl family kinases when the virus fuses with host plasma membrane during its egress (Newsome et al., 2004).

To ensure we examined the relevant population of A36, we directly looked at the level of Tyr132 phosphorylation associated with individual virions inducing actin tails using an antibody that detects pTyr132 (Newsome et al., 2004). We found that the antibody labels A36 N-G and G-N but not N-X (negative control lacking Grb2 binding site) virus particles inducing actin tails (Figure 6A). The lack of labelling of the N-X virus demonstrates that the antibody is specific as it does not detect any other phosphorylated component in the signalling network involved in nucleating actin tails. Moreover, the intensity of labelling was significantly higher on the G-N compared to the N-G virus, suggesting there is no impairment in phosphorylation. This enhanced labelling intensity most likely reflects increased antibody access to its pTyr132 epitope in the absence of Grb2 recruitment. To test this possibility, we determined the level of pTyr132 antibody labelling on the A36 N-G virus when Grb2 was knocked down using siRNA. Consistent with our hypothesis, we found that a reduction in levels of Grb2 results in increased labelling on the A36 N-G virus (p=0.071, Figure 6B). This increase was less dramatic than the G-N virus as the knockdown was not complete and the virus is good at recruiting any remaining Grb2. Taken together, our observations suggest that the lower levels of Grb2 recruitment observed when its binding site is repositioned are a consequence of changes in the signalling network rather than a reduction in phosphorylation of Tyr132.

**Figure 6:**
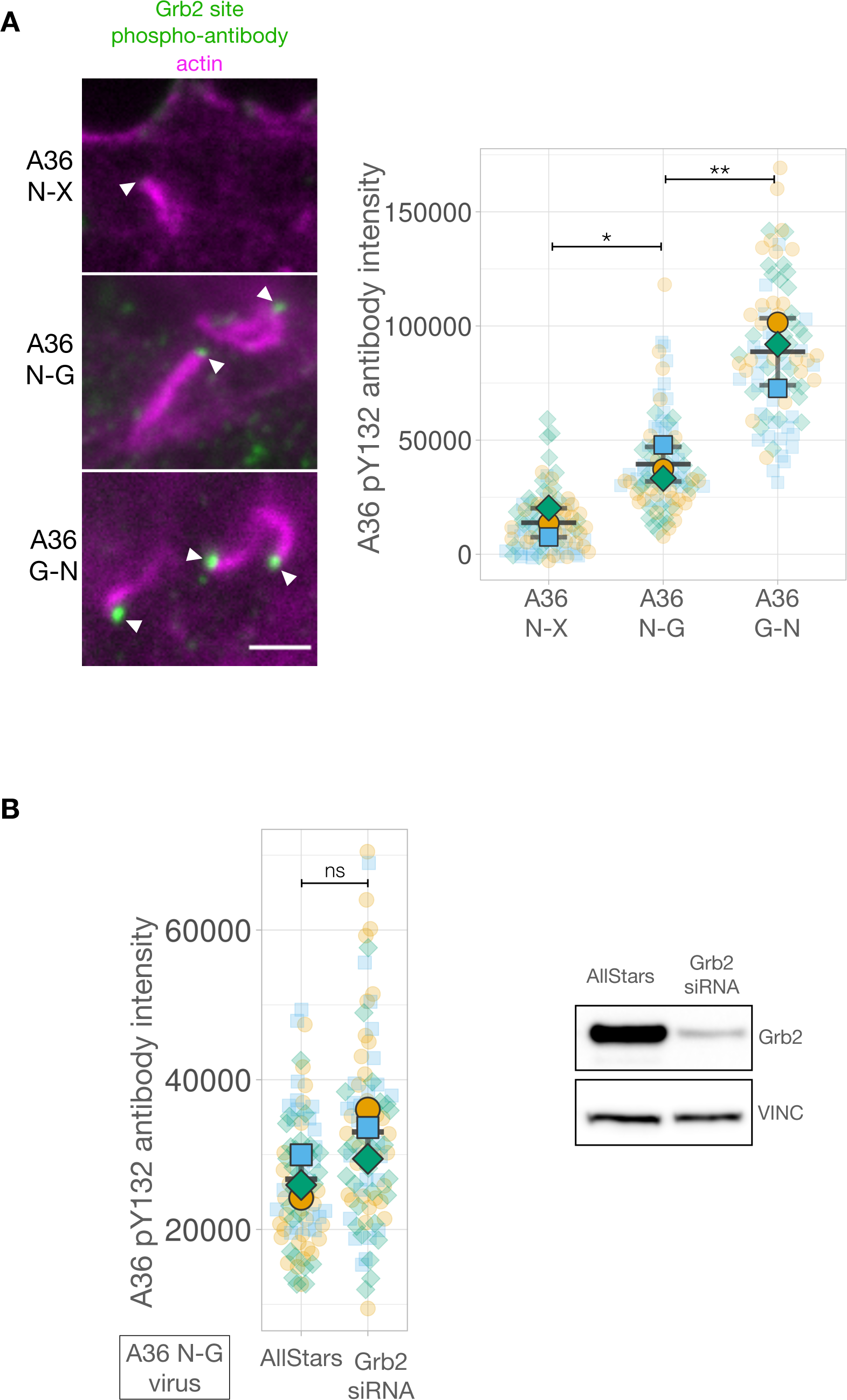
The Grb2-binding site in A36 is phosphorylated when motif positions are changed. **A.** Representative immunofluorescence images of A36 pY132 antibody labelling of indicated virus inducing actin tails in HeLa cells at 8 hours post-infection. Actin is stained with phalloidin. A36 N-X is a virus where the A36 Grb2-binding site is disrupted with by Tyr to Phe point mutation. The white arrowheads indicate virus position. Scale bar = 1 μm. The graph shows quantification of background-subtracted antibody intensity at the tip of actin tails. **B.** Quantification of background-subtracted A36 pY132 antibody intensity in HeLa cells infected with the A36 N-G virus and treated with indicated siRNA. The p value here is 0.071. On the right a representative immunoblot from one of three experimental repeats is shown. Vinculin levels were used as a loading control. Intensity of 85 virus particles was measured in three independent experiments. All error bars represent S.D. and the distribution of the data from each experiment is shown using a “SuperPlot”. Tukey’s multiple comparison’s test (for panel A) and Welch’s t test (for panel B) were used to determine statistical significance; ns, p >0.05; * p ≤ 0.05; ** p ≤ 0.01.

### Position rather than the number of Grb2 sites influences signalling network output

Our data indicate that the reduced signalling output of the A36 G-N virus is linked to decreased levels of Grb2 in the signalling network. This suggests that the position of Grb2 binding relative to Nck is the critical factor influencing the functional outcome of the network formed by the A36 G-N virus. We therefore wondered what impact an additional Grb2 site would have on virus-induced actin polymerization? More importantly, would any improvement in signalling output be dependent on the position of this second Grb2 site relative to Nck binding? To address this question, we generated recombinant G-N viruses expressing A36 with an extra Grb2 site N-terminal (A36 G-G-N virus) or C-terminal (A36 G-N-G virus) to the Nck site (Figure 7A). All four viruses (N-G, G-N, G-N-G and G-G-N) recruited similar levels of Nck and induced comparable numbers of actin tails (Figure 7 – supplement 1A and B). Nevertheless, the A36 G-N and G-G-N viruses induced the formation of equally short actin tails (Figure 7B). In contrast, the actin tails formed by the A36 G-N-G virus are noticeably longer. A similar trend was also observed for virus speed and spread (Figure 7C, D). Strikingly, the G-N-G and N-G viruses recruit similar levels of N-WASP, which are ∼ 2- fold greater than the G-G-N and G-N viruses (Figure 7E). They also recruit significantly more GFP-Grb2 (Figure 7 – supplement 1C). Grb2 is not essential for Vaccinia actin tail formation but its binding position relative to Nck clearly influences the level of N-WASP recruitment. This has a profound influence on the organization and output of the signalling network stimulated by phosphorylation of A36.

**Figure 7:**
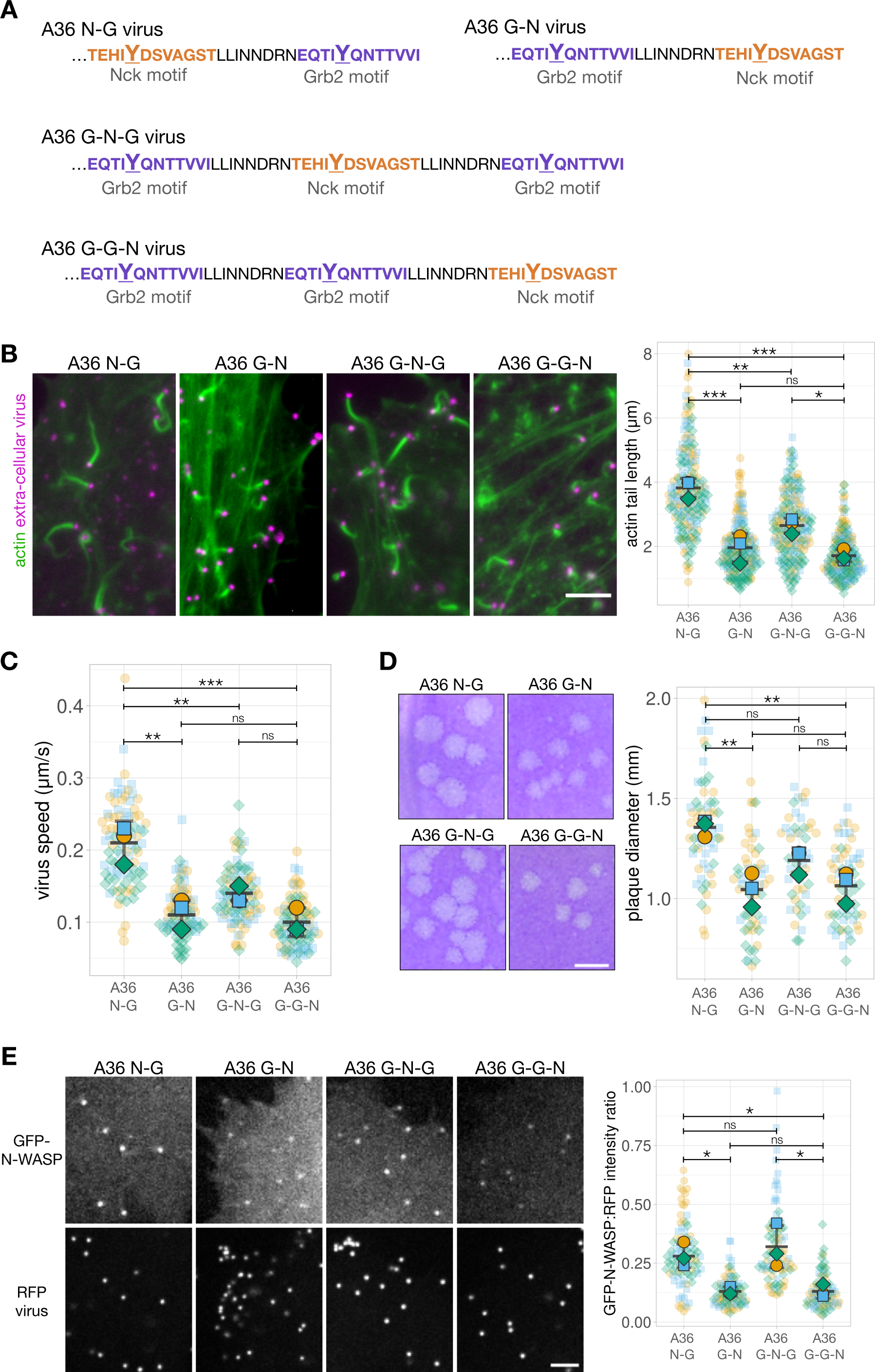
Signalling output can be improved by adding a new Grb2 binding site, but only in a position-dependent fashion. **A.** C-terminal amino acid sequence of A36 in recombinant viruses showing the position of phosphotyrosine motifs in wild type (A36 N-G) and swapped (A36 G-N) configurations and with a new Grb2-binding site C-terminal (G-N-G) or N-terminal (G-G-N) to the swapped configuration. **B.** Representative immunofluorescence images of actin tails in HeLa cells infected with the indicated virus at 8 hours post-infection. Actin is stained with phalloidin, and extra-cellular virus particles are labelled using an anti-B5 antibody. Scale bar = 5 μm. The graph shows quantification of actin tail length. 195 actin tails were measured in three independent experiments. **C.** The graph shows quantification of virus speed from time-lapse movies tracking the motility of the indicated RFP-A3-labelled virus over 50 seconds at 8 hours post infection. 90 virus particles were tracked in three independent experiments. **D.** Representative images and quantification of plaque diameter produced by the indicated virus in confluent BS-C-1 cells 72 hours post-infection. 60 plaques were measured in three independent experiments. Scale bar = 2 mm. **E.** Representative images showing GFP-N- WASP and RFP-A3 intensity on virus particles in live HeLa cells infected with the indicated viruses recorded 8 hours post-infection. Scale bar = 4 μm. GFP N-WASP intensity data for A36 N-G and G-N viruses is the same as in Figure 5C. The graph shows quantification of GFP- N-WASP:RFP-A3 fluorescence intensity ratio. Intensity of 90 virus particles was measured in three independent experiments. All error bars represent S.D and the distribution of the data from each experiment is shown using a “SuperPlot”. Tukey’s multiple comparison test was used to determine statistical significance; ns, p >0.05; * p≤0.05; ** p ≤ 0.01; *** p ≤0.001.

## DISCUSSION

The pTyr motifs in Vaccinia A36 engage the SH2-SH3 adaptors Nck and Grb2, which play essential roles in many different cellular signalling cascades. For example, Grb2 directly binds receptors EGFR and Ras via its SH2 domain, while the SH2 domain of Nck binds PDGFR and ephrinb1 receptors among others (Bong et al., 2003; Cowan and Henkemeyer, 2018; Lettau et al., 2009; Nishimura et al., 2005; Pramatarova et al., 2003). These adaptors also participate in signalling networks that can undergo phase transitions, including those assembled by the membrane proteins LAT and nephrin in T-cell activation and kidney podocytes respectively (Case et al., 2019; Ditlev et al., 2019; Kim et al., 2019; Pak et al., 2016; Su et al., 2016). LAT engages Grb2 for its function while nephrin relies on Nck. In the case of Vaccinia, the recruitment of Nck and Grb2 at the plasma membrane leads to the recruitment of a signalling network that induces the formation of Arp2/3-dependent actin polymerization that acts to enhance the spread of viral infection (Frischknecht et al., 1999; Ward and Moss, 2004). The virus provides a model system to dissect this signalling network in a true physiological context as the output is robust, can be easily quantified and manipulated using recombinant viruses and different cell lines (Scaplehorn et al., 2002; Weisswange et al., 2009).

We have taken advantage of this model to address whether the relative positioning of pTyr motifs plays a role in the output of the system, as this is an unexplored aspect of pTyr networks. To tackle this question, we used a truncated variant of A36, which still recruits both Nck and Grb2 to induce robust actin polymerization. In addition, we also generated a recombinant virus expressing the cytoplasmic C-terminus of the unrelated integral membrane p14 from Orthoreovirus. p14 promotes N-WASP and Arp2/3 dependent cell fusion during Orthoreovirus infection by interacting with Grb2 using a pTyr116 motif located in its short cytoplasmic tail (Chan et al., 2020). We chose p14 as Scansite and ELM predictions suggested that it may also contain a Nck binding site 20 residues upstream of the Grb2 binding site (Kumar et al., 2019; Obenauer et al., 2003). Using these two models, we have uncovered that the relative position of the Nck and Grb2 pTyr binding motifs has a profound impact on the output of the signalling network. In the wild type situation, the Nck binding site (pTyr112 and pTyr96 in A36 and p14 respectively) plays the dominant role in promoting actin polymerization, while Grb2 plays a supporting role. This is consistent with previous *in vitro* observations demonstrating that Grb2 is significantly less efficient than Nck at promoting actin polymerization via N-WASP and Arp2/3 (Carlier et al., 2000; Okrut et al., 2015). Swapping the position of the two adaptor binding sites severely impairs actin-based motility and spread of the virus. The impairment seen in the swapped configuration of A36 correlates with reduced N-WASP accumulation at the virus, even though the levels of Nck remain the same. The levels of Grb2 are also reduced, even though Tyr132 is still phosphorylated. By introducing a second Grb2 site N or C-terminal to Nck when it is in the “swapped” configuration, we found that the relative position of Grb2 has a dramatic impact on the system regardless of the level of Nck, which remains constant in our different recombinant viruses. The output of the network clearly does not merely depend on the level of Nck recruitment. This suggests that Grb2 places constraints on the organization of the network that limit or impede the ability of Nck to bind N-WASP and activate the Arp2/3 complex to promote actin polymerization.

So why does the position of Grb2 recruitment relative to that of Nck matter? Nck and Grb2 are multivalent adaptors that can interact with N-WASP to activate Arp2/3. Grb2 has two SH3 domains that bind N-WASP while the three SH3 domains of Nck engage with WIP and N-WASP (Carlier et al., 2000; Donnelly et al., 2013; Rivera et al., 2004). In addition, the interdomain linker of Nck also contributes to N-WASP activation (Banjade et al., 2015; Bywaters and Rivera, 2021; Okrut et al., 2015). One would envisage that these different interactions offer the possibility of activating N-WASP via a number of different routes. This notion would also be consistent with the concept of phase transition, which is dependent on stochastic multivalent interactions (Fuxreiter and Vendruscolo, 2021; Lyon et al., 2021). However, our observations clearly demonstrate that there is a preferred configuration or directionality to achieve the optimal network output. This implies that the network has underlying wiring principles.

Our finding that optimal signalling outputs are only achieved with specific pTyr motif configurations has important implications for membrane-associated complexes in the field of cellular condensates as well as in modular signalling. Recent evidence points to organizational principles that could govern the function of other cellular condensates such as P granules, stress granules and puncta formed by the FUS (fused in sarcoma) protein (Bienz, 2020; Folkmann et al., 2021; Jain et al., 2016; Kato and McKnight, 2017; Kato et al., 2012). But as of yet, no rules have been elucidated for membrane signalling. Our observations are not predicted by a phase transition-based model, where high conformational entropy and multiple binding configurations are assumed to be available (Fuxreiter and Vendruscolo, 2021; Lyon et al., 2021). This is thought to be the case particularly when signalling motifs are present in disordered regions of proteins. Disordered proteins, which constitute 40% of the human proteome, are rich in short linear motifs (SLiMs) (Tompa et al., 2014; Wright and Dyson, 2014). Due to their ubiquity and modularity, networks assembled via SLiMs in unstructured peptides are of immense interest to biologists building synthetic signalling systems (Lim, 2010). In synthetic networks, the optimal organisation of globular domains is recognized to influence functionality, for example in the construction of chimeric antigen receptors (Finney et al., 1998). However, the relative position of SLiMs within a complex network has never been considered to play a role. As viral proteins are enriched in disordered regions with short host-mimicking motifs (Davey et al., 2011; Uversky, 2019), they offer unique tools to explore the importance of SLiMs positioning on signalling output, given the implications for signalling networks undergoing phase transition. Our observations also demonstrate that Vaccinia provides an excellent model platform to dissect additional signalling networks (physiological or synthetic) activated by Src and Abl family kinases.

## METHODS

### Expression constructs and targeting vectors

The expression vectors pE/L-LifeAct-iRFP670 (Galloni et al., 2021), pLVX-GFP-N-WASP (Donnelly et al., 2013) and pBS SKII RFP-A3L targeting vector (Weisswange et al., 2009) were previously made in the Way lab. The lentiviral expression construct pLVX-GFP-Nck was generated by sub-cloning the Nck1 coding sequence (Donnelly et al., 2013) into NotI/EcoRI sites of a pLVX-N-term-GFP parent vector (Abella et al., 2016). All other expression constructs generated for this study were made using Gibson Assembly (New England Biolabs) according to manufacturer’s instructions. Desired amino acid substitutions were introduced by whole-plasmid mutagenesis using complementary mutagenic primers. All primers used in cloning are listed in Table 1. The lentiviral expression construct pLVX-GFP- Grb2 was generated by cloning a GFP-Grb2 (Weisswange et al., 2009) fragment into the XhoI/EcoRI sites of a pLVX parent vector (Abella et al., 2016). The A36R-targeting vector was generated by amplifying a fragment containing the A36R gene including 325 bp upstream and downstream sequences from the WR strain of Vaccinia virus genomic DNA and cloning into the NotI/HindIII sites of pBS SKII. This vector was modified to generate desired A36 truncations and variants. The tagGFP2 coding sequence used in the A36 N-G and A36-G-N fusion constructs was amplified from a plasmid provided by David Drubin (UC Berkeley) (Akamatsu et al., 2020). The A36-p14 chimeric construct was obtained as a synthetic gene (Invitrogen; Geneart) and cloned into the A36R-targeting vector using SpeI/BsrGI sites in the sequences flanking the A36R coding region. SnapGene software (from Insightful Science; available at snapgene.com) was used to plan and visualise cloning strategies, and to analyse sequencing results.

**Table 1.**
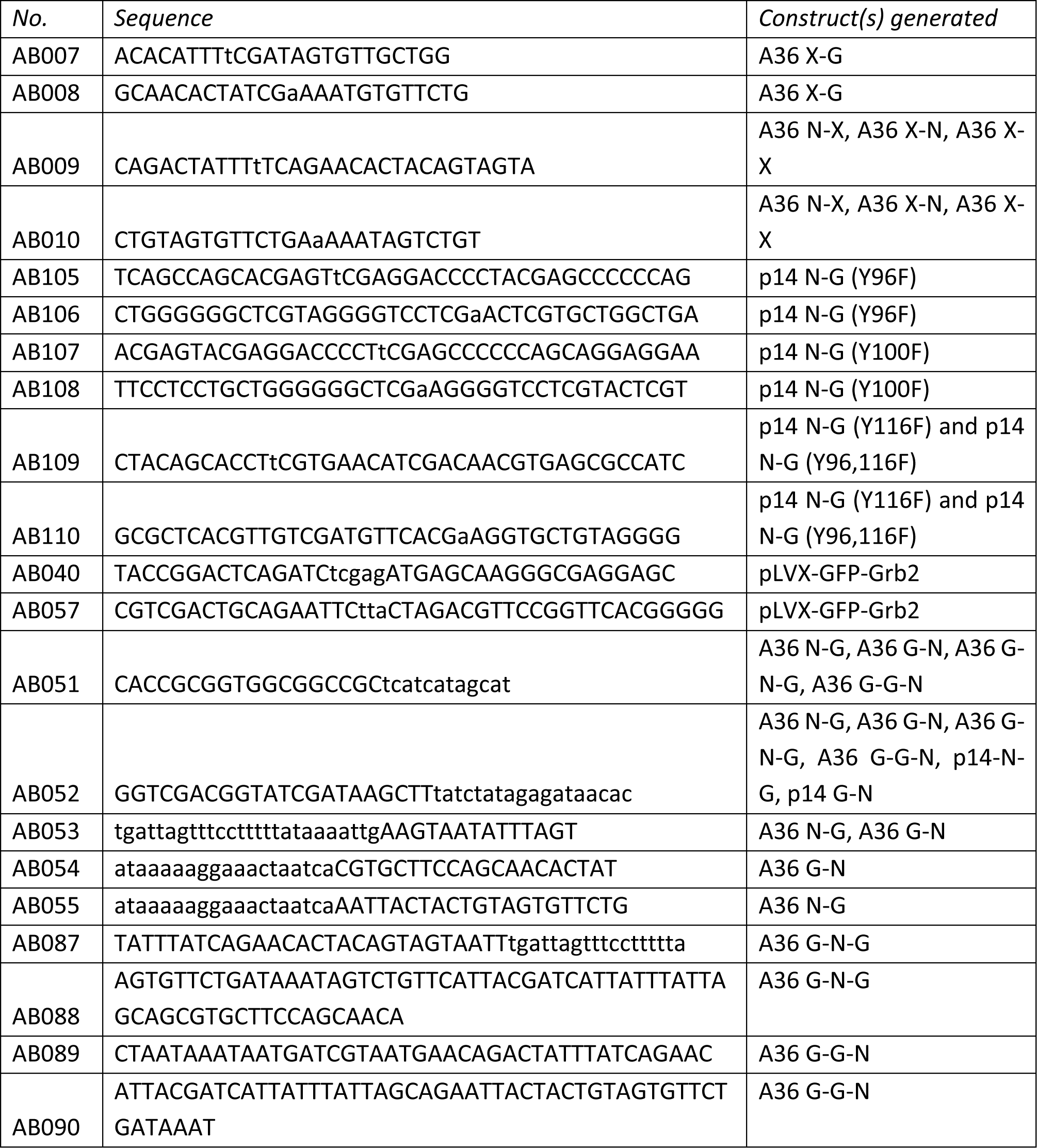
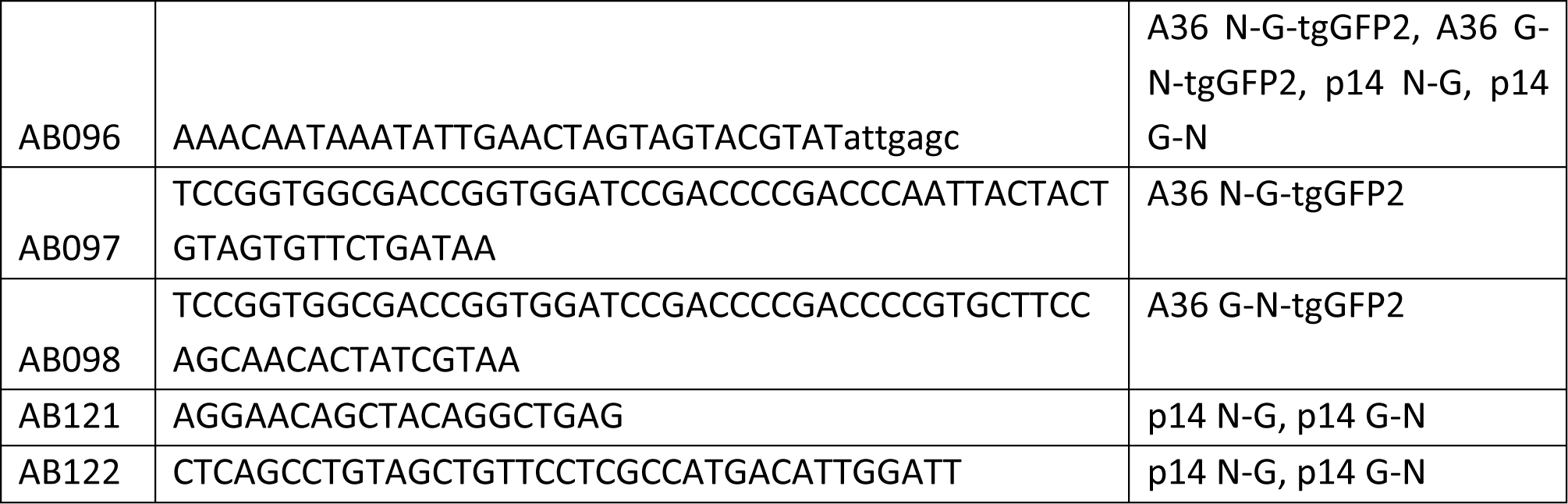
Primers

### Cell lines

All cell lines were maintained in minimal essential medium (MEM) supplemented with 10% FBS, 100 U/ml penicillin, and 100 µg/ml streptomycin at 37°C and 5% CO2. HeLa cell lines stably expressing LifeAct-iRFP670 (Snetkov et al., 2016) and GFP-WIP (Weisswange et al., 2009) were previously generated in the Way lab. Nck -/- MEFs (Bladt et al., 2003) and N- WASP-/-MEFs (Snapper et al., 2001) were provided by the late Tony Pawson (Samuel Lunenfeld Research Institute, Toronto, Canada) and Scott Snapper (Harvard Medical School, Boston, MA) respectively. For this study, lentiviral expression vectors were used to stably express GFP-Nck in HeLa cells and Nck-/- MEFs, GFP-N-WASP in HeLa cells and N-WASP-/- MEFs, and GFP-Grb2 in HeLa cells. All cell lines were generated using the lentivirus Trono group second generation packaging system (Addgene) and selected using puromycin resistance (1 µg/ml) as previously described (Abella et al., 2016). Expression of the relevant fusion proteins was confirmed by live imaging and immunoblot analysis (Suppl. Figure 5). The following primary antibodies were used: anti-Nck (BD transduction; 1:1000), anti-vinculin (Sigma #V4505; 1:2000), anti-Grb2 (BD Transduction #610112, 1:3000), anti-N-WASP (Cell Signalling #4848S; 1:1000), anti-GFP (3E1 custom made by Cancer Research UK; 1:1000). HRP-conjugated secondary antibodies were purchased from The Jackson Laboratory.

### Viral plaque assays

Plaque assays were performed in confluent BS-C-1 cell monolayers. Cells were infected with the relevant Vaccinia virus at a multiplicity of infection (MOI) = 0.1 in serum-free MEM for one hour. The inoculum was replaced with a semi-solid overlay consisting of a 1:1 mix of MEM and 2% carboxymethyl cellulose. Cells were fixed with 3% formaldehyde at 72 hours post-infection and subsequently visualized with crystal violet cell stain as previously described (Humphries et al., 2012). To determine plaque size, the diameter of well-separated plaques was measured using the Fiji line tool (Schindelin et al., 2012).

### Construction of recombinant Vaccinia viruses

In this study, recombinant Vaccinia viruses in the WR background were isolated by selecting viral plaques based on their size or by introduction of a fluorescently-tagged viral protein as described previously (Snetkov et al., 2016; Weisswange et al., 2009). The former strategy was used to generate recombinants where A36 variants were introduced at the endogenous locus by rescuing plaque size in the WR–ΔA36R virus that makes very small plaques (Parkinson and Smith, 1994; Ward et al., 2003). To introduce relevant constructs into the A36 genomic locus, HeLa cells infected with WR–ΔA36R at MOI = 0.05 were transfected with the appropriate pBS SKII A36R-targeting vectors using Lipofectamine2000 (Invitrogen) as described by the manufacturer. In the case of the p14 N-G virus, a PCR fragment containing the desired construct flanked by recombination arms was used for transfection. When all cells displayed cytopathic effect at 48-72 hours post-infection, they were lysed, and serial dilutions of the lysates were used to infect confluent BS-C-1 cell monolayers in a plaque assay (see above). Plaques were revealed by neutral red staining and recombinants were identified and picked based on increased plaque size. Plaque lysates were used to infect fresh BS-C-1 cell monolayers over at least three rounds of plaque purification. To isolate A36 N-G-tgGFP2 and A36 G-N-tgGFP2 viruses, in addition to size, plaque fluorescence was used to identify recombinants. To generate recombinants where the viral core was fluorescently labelled with RFP, HeLa cells infected with the relevant parent virus were transfected with the pBS SKII RFP-A3-targeting vector (Weisswange et al., 2009).

Recombinant viruses were isolated based on RFP fluorescence over at least three rounds of plaque purification. In all cases, successful recombination at the correct locus, loss of the parent variant and virus purity were verified by PCR and sequencing. All recombinant viruses were purified through a sucrose cushion before use and storage.

### Vaccinia virus infection for imaging

For live and fixed cell imaging, cells were infected with the relevant Vaccinia virus recombinant in serum-free MEM at MOI = 1. After one hour at 37°C, the serum-free MEM was removed and replaced with complete MEM. Cells were incubated at 37°C until further processing.

### Transient transfection and siRNA

Transient transfection of A36-p14 hybrid constructs and the pE/L-LifeAct-iRFP670 expression in Vaccinia-infected cells was done using FUGENE (Promega) as described by the manufacturer. To transiently express the A36-p14 chimera and its variants (Figure 3B to E), expression vectors containing the relevant construct under the control of the A36 promoter were transfected into cells one hour after infection with the WR-ΔA36 virus (Parkinson and Smith, 1994). To visualize actin structures in live HeLa cells stably expressing GFP-Grb2 (Figure 4A), cells were transfected with pE/L-LifeAct-iRFP670 (Galloni et al., 2021) one hour after infection with p14 N-G virus. For knockdown experiments, HeLa cells were transfected with siRNA as previously described (Abella et al., 2016). Cells were infected with Vaccinia virus 72 hours after siRNA transfection, and samples from each siRNA condition were kept for immunoblot analysis. The following siRNAs were used: AllStars (Qiagen; SI03650318), Grb2-targeting siRNA oligos GAAAGGAGCUUGCCACGGGUU and CGAAGAAUGUGAUCAGAACUU. The following primary antibodies were used in immunoblots: anti-vinculin (Sigma #V4505; 1:2000), anti-Grb2 (BD Transduction #610112, 1:3000). HRP-conjugated secondary antibodies were purchased from The Jackson Laboratory.

### Immunofluorescence

At 8 hours (HeLa) or 16 hours (MEFs) post-infection, cells were fixed with 4% paraformaldehyde in PBS for 10 min, blocked in cytoskeletal buffer (1 mM MES, 15 mM NaCl, 0.5 mM EGTA, 0.5 mM MgCl2, and 0.5 mM glucose, pH 6.1) containing 2% (vol/vol) fetal calf serum and 1% (wt/vol) BSA for 30 min, and then permeabilized with 0.1% Triton-X/PBS for 5 min. To visualize cell-associated enveloped virions (CEV), cells were stained with, with a monoclonal antibody against B5 (19C2, rat, 1:1000; (Hiller and Weber, 1985)) followed by an Alexa Fluor 647 anti-rat secondary antibody (Invitrogen; 1:1000 in PBS) prior to permeabilization of the cells with detergent. Other primary antibodies used were anti-Nck (Millipore #06-288; 1:100), anti-WIP (1:100; (Moreau et al., 2000)), and anti-N-WASP (Cell Signalling #4848S; 1:100) followed by Alexa Fluor 488 conjugated secondary antibodies (Invitrogen; 1:1000 in PBS). Actin tails were labeled with Alexa Fluor 488, Alexa Fluor 568 or Alexa Fluor 647 phalloidin (Invitrogen; 1:500). Coverslips were mounted on glass slides using Mowiol (Sigma). Coverslips were imaged on a Zeiss Axioplan2 microscope equipped with a 63x/1.4 NA Plan-Achromat objective and a Photometrics Cool Snap HQ cooled charge-coupled device camera. The microscope was controlled with MetaMorph 7.8.13.0 software. To measure the levels of pY132 at the virus (Figure 6) coverslips were fixed with 4% paraformaldehyde containing 0.1% Triton-X prior to staining with an antibody against the A36 phosphotyrosine 132 site (1:100, (Newsome et al., 2004)). Mounted coverslips were imaged on an Olympus iX83 Microscope with Olympus 100x/1.50NA A-Line Apochromatic Objective Lens, dual Photometrics BSI-Express sCMOS cameras and CoolLED pE-300 Light Source. The microscope was controlled with MicroManager 2.0.0 software.

### Live-cell imaging

Live-cell imaging experiments were performed at 8 hours (HeLa) or 16 hours (MEFs) post-infection in complete MEM (10% FBS) in a temperature-controlled chamber at 37°C. Cells were imaged on a Zeiss Axio Observer spinning-disk microscope equipped with a Plan Achromat 63x/1.4 Ph3 M27 oil lens, an Evolve 512 camera, and a Yokagawa CSUX spinning disk (Galloni et al., 2021; Pfanzelter et al., 2018). The microscope was controlled by the SlideBook software (3i Intelligent Imaging Innovations). For determining recruitment levels of GFP-tagged molecules to the virus, single snapshots of live cells were acquired. To determine virus speed, images were acquired for 50 seconds at 1Hz.

### Image analysis and quantitation

Quantification of actin tail number and length was performed using two-colour fixed cell images where actin and extracellular virus were labelled. Ten cells were analysed per condition in each independent experiment. The number of actin tails was measured by randomly selecting 25 isolated extracellular virus particles in each image and determining the presence of a tail in the corresponding actin channel. Actin tail length of 8 randomly selected tails per image was measured using the freehand line drawing function in Fiji. To analyse virus motility, two-colour time-lapse movies of HeLa cells stably expressing LifeAct- iRFP670 infected with the relevant recombinant virus labelled with RFP-A3 were used. The velocity of virus particles in the RFP channel was measured using a Fiji plugin developed by David Barry (the Francis Crick Institute) as previously described (Abella et al., 2016). Bona fide actin-based virus motility was verified manually using the corresponding iRFP670 channel. Five movies were analysed per condition, and speeds from 30 particles were measured in each independent experiment. pY132 antibody intensity was analysed in two-colour fixed cell images where actin was co-labelled. Raw integrated density of the antibody signal was measured at the tip of actin tails after local background subtraction using Fiji. Thirty particles were measured per condition in each independent experiment. Recruitment levels of GFP-tagged molecules to the virus was analysed in two-colour live cell images as an intensity ratio to RFP-A3. Cells with many virus particles at the cell periphery were selected for analysis. GFP images were background subtracted using a median filtered image. The ratio of GFP:RFP intensity at peripheral particles was then measured using Fiji. Five images were analysed per condition, and 30 particles were measured in each independent experiment. In Figure 5B, GFP-Nck intensity at extracellular virus particles was measured in fixed cells using a Fiji particle mapping plugin (Wanaguru et al., 2018).

### Phosphopeptide pulldown assay

Phosphorylated and non-phosphorylated peptides were synthesised in-house (p14: SQHE<u>pY</u>EDPYEPP, SQHEYEDPYEPP; A36: APSTEHI<u>pY</u>DSVAGST, APSTEHIYDSVAGST) containing the relevant Nck-binding sites and an N-terminal biotin tag. These were coupled to streptavidin Dynabeads M-280 (Thermo Fischer Scientific). Uninfected HeLa cells were lysed in a buffer containing 50mM Tris.HCl pH7.5, 150mM NaCl, 0.5mM EDTA, 0.5% NP40, 0.5% Triton-X and a cocktail of protease and phosphatase inhibitors (1mM orthovanadate, cOmplete (Roche), PHOSstop (Roche)). A postnuclear supernatant was obtained by a 16,000g centrifugation for 10 min at 4°C. The peptide-coupled beads were incubated with these clarified cell lysates. Unbound proteins were removed from beads in three washes in the same cell lysis buffer. The proteins bound to beads were resolved on an SDS-PAGE and the presence of Nck was determined by immunoblot analysis using anti-Nck antibody (Millipore #06-288; 1:1000). As a negative control anti-Grb2 (BD Transduction #610112, 1:3000) was used. HRP-conjugated secondary antibodies were purchased from The Jackson Laboratory.

### Statistical analysis and figure preparation

All data are presented as means ± S.D. For all experiments, means of at least three independent experiments were used to determine statistical significance by a Welch’s t-test (comparing only two conditions), Tukey’s multiple comparisons test (comparing multiple conditions with each other) or a Dunnett’s multiple comparisons test (comparing multiple conditions with a control). All data are represented as SuperPlots to allow assessment of the data distribution in individual experiments (Lord et al., 2020). SuperPlots were generated using the SuperPlotsOfData webapp (Goedhart, 2021) and graphs showing intrinsic disorder predictions were generated in GraphPad Prism 9. All data were analyzed using GraphPad Prism 9 or the SuperPlotsOfData webapp. Temporal overlays of live imaging data to illustrate virus motility were generated using the temporal colour-code function in Fiji. Schematics were created with BioRender.com. Final figures were assembled using Keynote software.

## ACKNOWLEDGEMENTS

We thank Nicola O’Reilly, Dhira Joshi and Stefania Federico (Peptide Chemistry, the Francis Crick Institute) for synthesizing peptides. We thank Cell Services and Genomics Equipment Park at the Francis Crick Institute for their help with maintaining cell lines and DNA sequencing respectively. We thank members of the Way Laboratory for useful discussions and suggestions, in particular Davide Carra, for nucleating the idea of adding an extra Grb2 site. We also thank Frank Uhlmann and Neil McDonald (the Francis Crick Institute) for helpful comments on the manuscript. Michael Way was supported by Cancer Research UK (FC001209), UK Medical Research Council (FC001209), and Wellcome Trust (FC001209) funding at the Francis Crick Institute. For the purpose of Open Access, the authors have applied a CC BY public copyright license to any Author Accepted Manuscript version arising from this submission.

## AUTHOR CONTRIBUTIONS

A.B. and M.W. designed the study. A.B. generated the required reagents and performed all experiments and analyses. A.B. and M.W. wrote the manuscript.

---

**Figure 1 - supplement 1:**
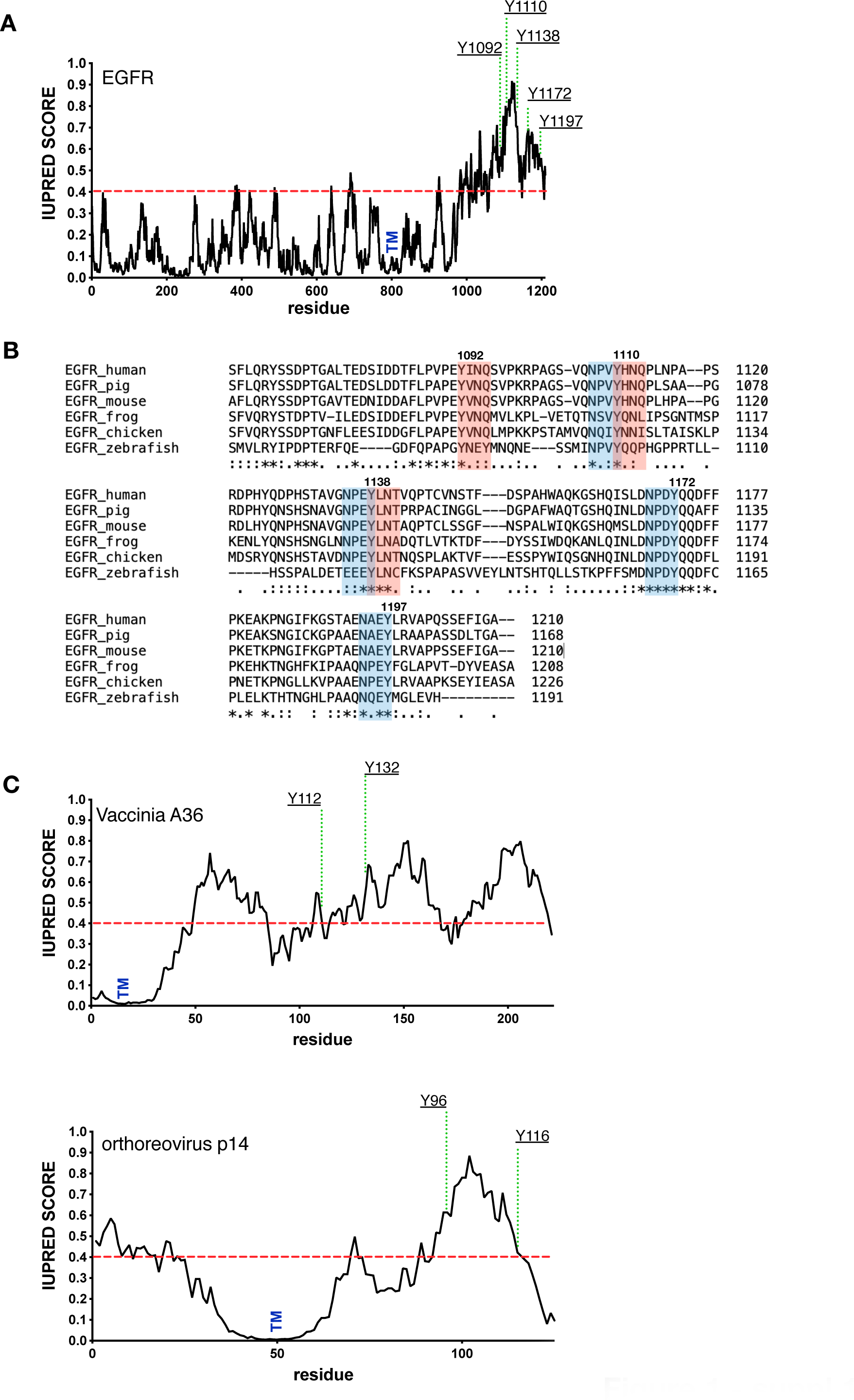
pTyr motifs are found in disordered regions. **A.** Shows a graph predicting the intrinsic disorder of human EGFR (https://iupred2a.elte.hu/). The positions of key phosphotyrosine residues and the transmembrane domain (TM) are indicated. **B.** Clustal sequence alignment of the C- terminus of EGFR from the indicated species. Positions of the conserved Grb2 and Shc1 adaptor binding sites are shown in pink and blue respectively. **C.** Shows the predicted intrinsic disorder of Vaccinia A36 and Orthoreovirus p14 together with the positions of key phosphotyrosine residues involved in recruiting Nck and Grb2 and the transmembrane domain (TM).

**Figure 2 - supplement 1:**
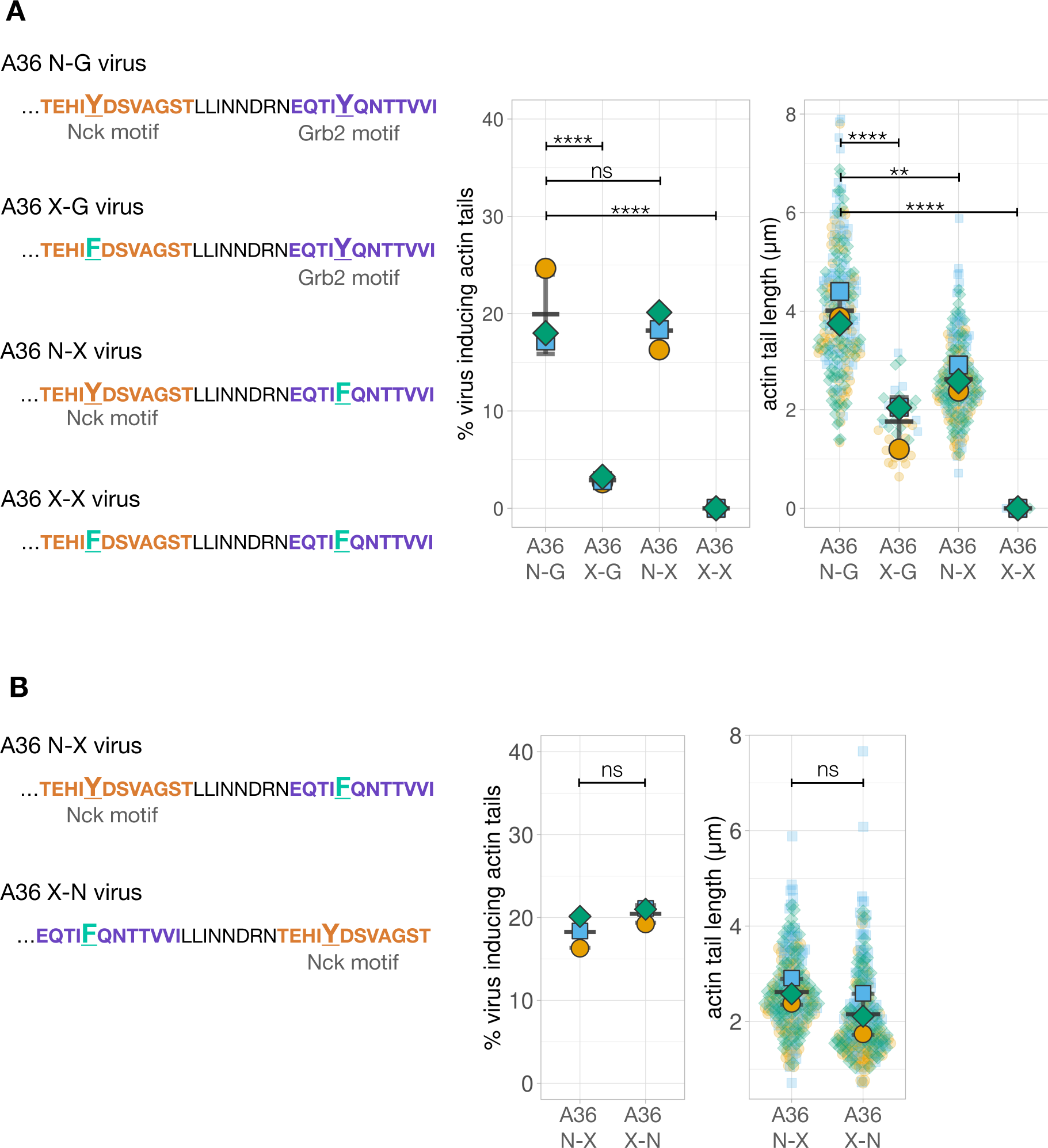
Quantification of the number of virus inducing actin tails along with their length. **A.** Left - Amino acid sequences of the indicated truncated A36 recombinants where either one or both phosphotyrosine motifs (orange for Nck and purple for Grb2) are disrupted with a Tyr (Y) to Phe (F) point mutation. Right - Quantification of the number of extracellular virus inducing actin tails together with their length. **B.** C-terminal amino acid sequences of truncated A36 recombinants with wildtype or swapped Nck binding sites but deficient in Grb2 recruitment (Phe substitution) together with quantification of the number of extracellular virus inducing actin tails together with their length. The same A36 N-X data is shown in **A** and **B**. In both cases, 240 actin tails were measured in three independent experiments, except in the A36 X-G mutant where very few actin tails were made. All error bars represent S.D. and the distribution of the data from each experiment is shown using a “SuperPlot”. Dunnett’s multiple comparison’s test was used to determine statistical significance; ns, p >0.05; ** p ≤ 0.01; **** p ≤0.0001.

**Figure 4 – supplement 1:**
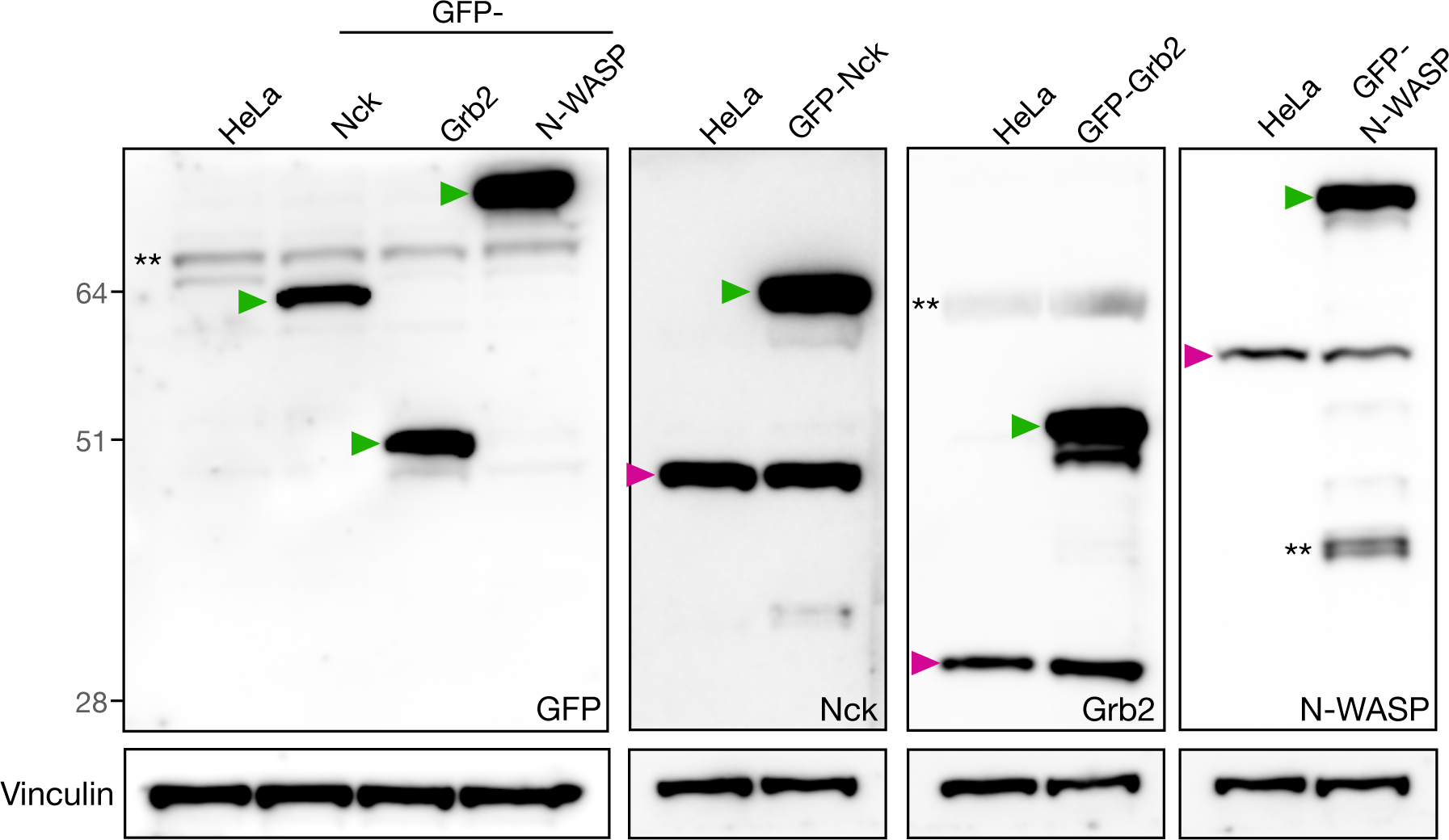
Validation of stable cell lines by immunoblot. Immunoblot analyses of total cell lysates from HeLa cell lines stably expressing GFP-Nck, GFP-Grb2 or GFP-N-WASP with the indicated antibodies. The green and magenta triangles indicate the GFP-tagged and endogenous proteins respectively. ** denotes non-specific bands.

**Figure 5 - supplement 1:**
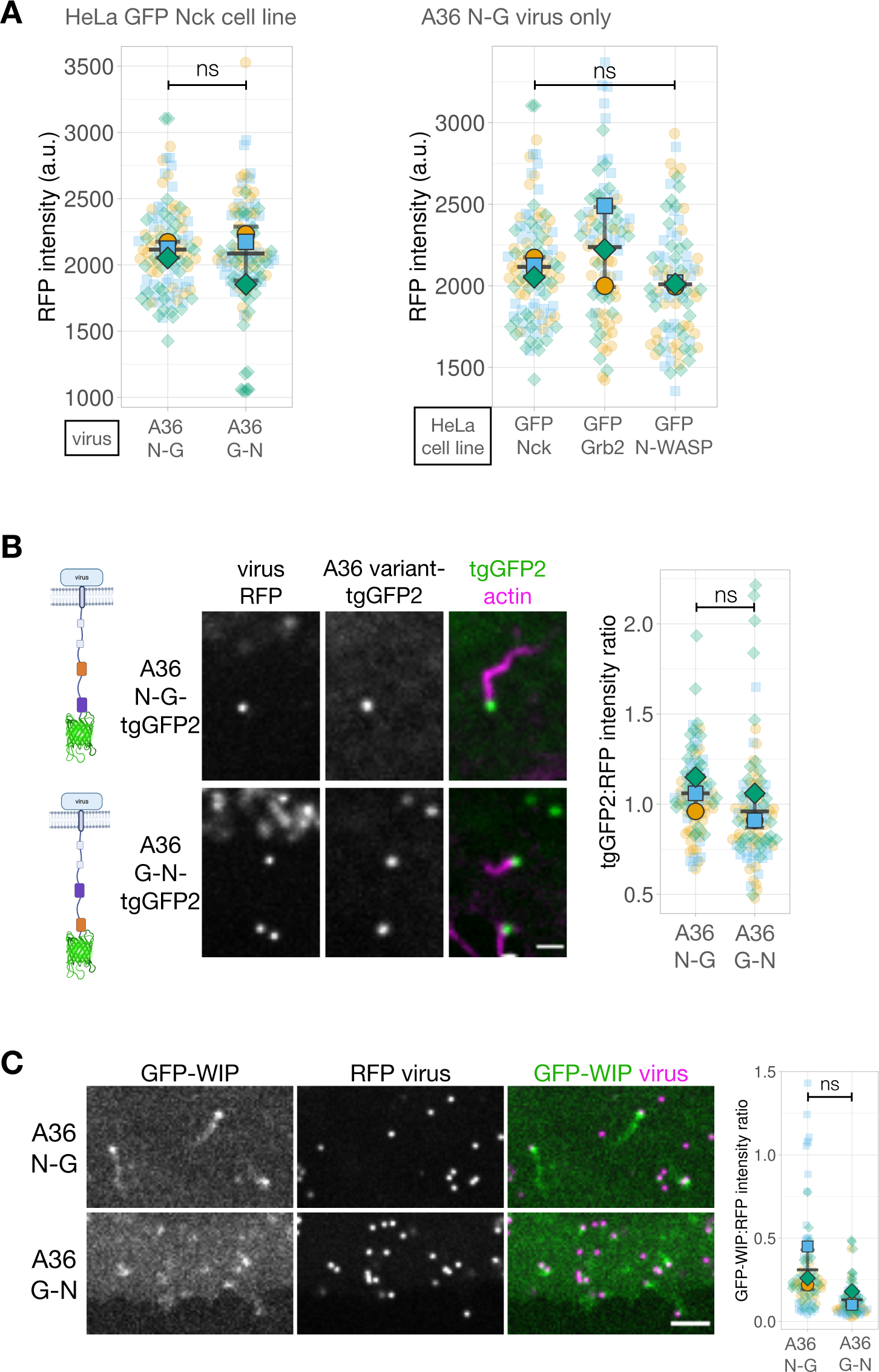
Fluorescence intensity measurements of Vaccinia recombinants. **A.** The graphs show quantification of RFP-A3 intensity on virus particles in different recombinants (left) and HeLa cell lines (right) measured in live cells at 8 hours post- infection. These datasets correspond to Figure 5A. Intensity of 90 virus particles was measured in three independent experiments. **B.** Left - A schematic showing the A36 variant expressed in each recombinant virus. The Nck-binding motif is indicated in orange, the Grb2-binding motif in purple and tgGFP2 in green. Middle - Representative images of tgGFP2 and RFP-A3 labeled virus particles in live HeLa cells stably expressing LifeAct iRFP infected with the indicated viruses recorded 8 hours post-infection. All virus particles are marked with an RFP tag fused to a viral core protein A3. Scale bar = 2 μm. Right - The graph shows quantification tgGFP2:RFP fluorescence intensity ratios at the tip of actin tails. Intensity of 95 virus particles was measured in three independent experiments. **C.** Representative images showing GFP-WIP recruitment to RFP-A3 labelled virus particles in live HeLa cells infected with the indicated viruses recorded 8 hours post-infection. Scale bar = 2 μm. The graph shows quantification of GFP-WIP:RFP-A3 fluorescence intensity ratio. Intensity of 90 virus particles was measured in three independent experiments. All error bars represent S.D and the distribution of the data from each experiment is shown using a “SuperPlot”. Dunnett’s multiple comparison’s test (for panel A right) or Welch’s t test (remaining panels) were used to determine statistical significance; ns, p >0.05; ** p ≤ 0.01.

**Figure 5 – supplement 2:**
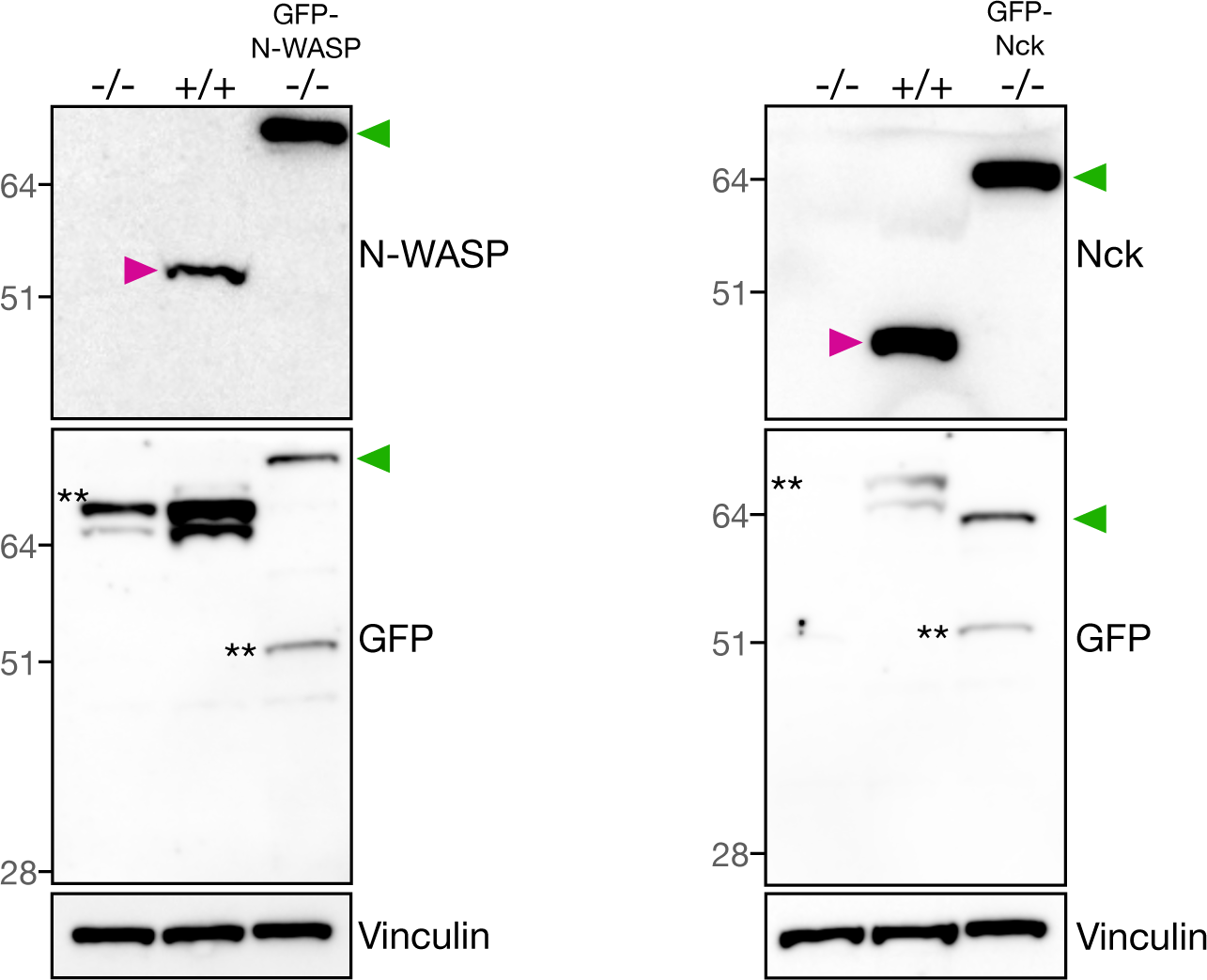
Verification of stable cell lines by immunoblot. Immunoblot analyses of total cell lysates from N-WASP or Nck null MEFs with or without stable expression of their respective GFP-tagged version together with their parental wildtype cells. Immunoblot analyses was performed with the indicated antibodies. The green and magenta triangles indicate the GFP-tagged and endogenous proteins respectively. ** denotes non-specific bands.

**Figure 7 - supplement 1:**
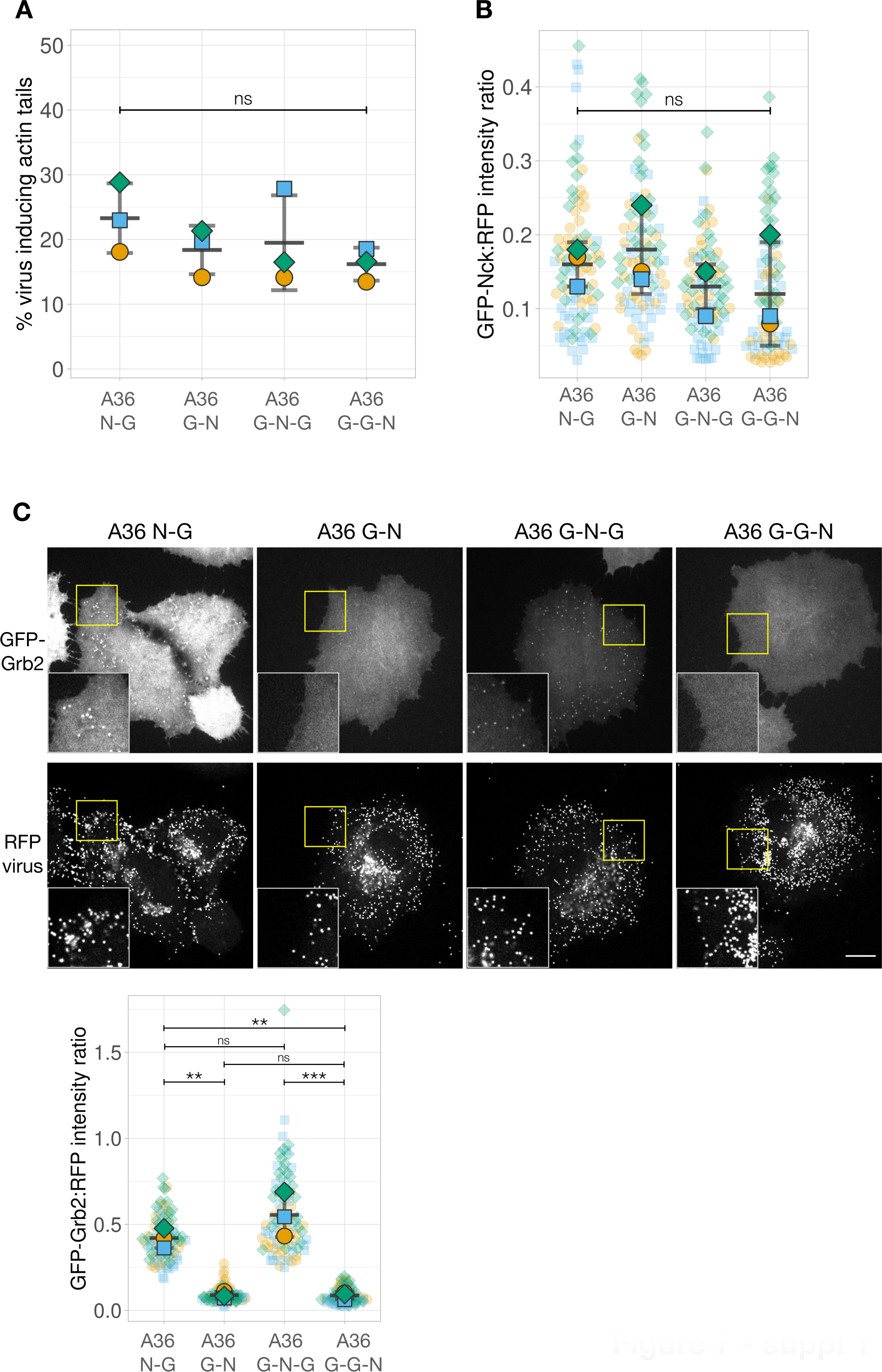
Quantification of the number of virus inducing actin tails and GFP-Nck recruitment in A36 variants with an extra Grb2-binding site. **A.** Quantification of number of extracellular virus particles inducing actin tails in the indicated viruses. **B.** Quantification of the GFP-Nck:RFP-A3 fluorescence intensity ratio of virus particles in live HeLa cells infected with the indicated viruses at 8 hours post-infection. The data for A36 N-G and G-N viruses is the same as in Figure 5C. Intensity of 85 virus particles was measured in three independent experiments. **C.** Representative images showing GFP-Grb2 and RFP-A3 intensity on virus particles in live HeLa cells infected with the indicated viruses recorded 8 hours post-infection. The insets show a 2x magnification of the yellow boxed regions. Scale bar = 16 μm. The graph shows quantification of GFP-Grb2:RFP- A3 fluorescence intensity ratio. Intensity of 90 virus particles was measured in three independent experiments. All error bars represent S.D. and the distribution of the data from each experiment is shown using a “SuperPlot”. Tukey’s multiple comparison test was used to determine statistical significance; ns, p >0.05; ** p ≤ 0.01; *** p ≤0.001.

**Supplemental Movie 1: Phosphotyrosine motif position impacts actin-based motility of Vaccinia virus.** HeLa cells stably expressing LifeAct-iRFP670 (green) were infected with either the A36 N-G or A36 G-N virus labelled with RFP-A3 (magenta) for 8 hours. Images were taken every second. Video plays at 5 frames per second. The RFP-A3 signal was used to generate the temporal colour-coded representation in Figure 2C. The time in seconds is indicated, and the scale bar = 3 µm.

**Supplemental Movie 2: Phosphotyrosine motif position impacts actin-based motility driven by p14 N-G virus.** HeLa cells stably expressing LifeAct-iRFP670 (green) were infected with either the p14 N-G or p14 G-N virus labelled with RFP-A3 (magenta) for 8 hours. Images were taken every second. Video plays at 5 frames per second. The RFP-A3 signal was used to generate the temporal colour-coded representation in Figure 4D. The time in seconds is indicated, and the scale bar = 3 µm.

## Notes

### Competing Interest Statement

The authors have declared no competing interest.

